# Multilingual Computational Models Capture a Shared Meaning Component in Brain Responses across 21 Languages

**DOI:** 10.1101/2025.02.01.636044

**Authors:** Andrea Gregor de Varda, Saima Malik-Moraleda, Greta Tuckute, Evelina Fedorenko

## Abstract

At the heart of language neuroscience lies a fundamental question: How does the brain process the rich variety of languages? Multilingual neural network models offer a way to answer this question by representing linguistic content across languages in a shared space. Leveraging these advances, we evaluated the similarity of linguistic representations in speakers of 21 languages. We combined existing (12 languages across 4 language families) and newly collected fMRI data (9 languages across 4 families) to test encoding models predicting brain activity in the language network using representations from multilingual models. Model representations reliably predicted brain responses within each language. Critically, encoding models can be transferred zero-shot across languages, so that a model trained to predict brain activity in a set of languages can account for responses in a held-out language. These results imply a shared cross-lingual component, which appears to be related to a shared meaning space.

## Introduction

Natural languages vary remarkably in how they convey meaning; this variety spans all levels of linguistic structure, from phonology to syntax (Evans & Levinson, 2009). Consider, for instance, the intricate verb conjugations in Italian, which convey tense, aspect, gender, and number, contrasting sharply with the more context-reliant structures in Mandarin, where verb forms are not inflected. In Finnish, the extensive case system can express complex structural relationships between nouns and other sentence elements, a feature largely absent in English. And the tonal modulations of Vietnamese create meanings distinguished by pitch, a concept foreign to non-tonal languages. Despite this variability, all human languages allow their speakers to express almost any idea in a sequence of words—describing objects, events, relations, and inner states. This expressive power makes human language the most sophisticated communication system in the animal kingdom. Here, we test whether the human language network—a set of brain areas that support human linguistic ability (Fedorenko et al., 2024a)—encodes linguistic inputs in a consistent way across diverse languages.

A core challenge in addressing this question is to quantitatively capture linguistic information consistently across languages. Recent progress in multilingual Natural Language Processing (NLP) research has provided a promising approach to represent linguistic content in a language-general format, making it possible to project words and sentences onto comparable representation spaces across languages. Such advances have been driven by the introduction of Multilingual Neural Network Language Models (MNNLMs). MNNLMs are transformer-based language models trained simultaneously on text in several languages (up to 200 languages in the case of NLLB-200; NLLB Team, 2024). Some of these models, such as machine translation models (NLLB Team, 2024), are provided with explicit cross-lingual supervision during training, so that their training data comprises translation-equivalent sentences that facilitate cross-lingual alignment. For other models (e.g., mBERT; Devlin et al., 2019), cross-lingual correspondences are not explicitly provided in the training data, but are instead implicitly learned from the statistical regularities across languages. These different models have shown remarkable ability in learning language-independent representation spaces, suggesting potential universality in linguistic feature encoding, including semantic and syntactic information (Chi et al., 2020; de Varda & Marelli, 2023, 2024; Stanczak et al., 2022). An appeal of these language-general linguistic representations is that they are constructed in a data-driven way without any potential bias that may be introduced by a particular linguistic theory or framework. To quantitatively capture linguistic content across languages, our study employs contextualized word and sentence embeddings derived from MNNLMs as representational spaces to understand the brain’s processing of linguistic inputs in diverse languages.

Several studies have now demonstrated that representations from neural network language models can successfully account for brain responses in the human language network (Antonello & Huth, 2024; Aw & Toneva, 2023; Caucheteux & King, 2022; Goldstein et al., 2022; Schrimpf et al., 2021; Tuckute et al., 2024a, 2024b, *inter alia*). These findings have been taken as evidence of the sensitivity of the human language processor to the distributional statistics of language and its engagement in predictive processing. However—similar to other work in the cognitive neuroscience of language (see Malik-Moraleda, Ayyash et al., 2022 for discussion; also, Blasi et al., 2022; Forgey, 2024)—these studies have been mostly conducted in English, with a few exceptions involving Dutch, French, Japanese, and Chinese (Caucheteux & King, 2022; Millet et al., 2022; Nakagi et al., 2024; Sun et al., 2023; Yamashita et al., 2023). Here, we first tested whether the relationship between language model representations and brain responses holds for diverse languages, thus plausibly reflecting key computational principles of human language processing. And then critically, we investigated whether representations in the language network are similar across languages.

A number of prior studies have shown that broad properties of the language network—including its left-lateralized topography and functional selectivity for language—are similar across languages and language families (Illes et al., 1999; Chee et al., 1999; Malik-Moraleda, Ayyash, et al., 2022). Some studies have gone beyond these broad cross-linguistic similarities, zooming in on the representations of linguistic content (Buchweitz et al., 2012; Correia et al., 2014, 2015; Sheikh et al., 2019, 2021; Zinszer et al., 2016). However, such studies have either examined the processing of multiple languages within bilingual speakers, where the similarity may reflect co-activation of the translation equivalents, especially in tasks that use single-word stimuli; or they used approaches, such as multivoxel pattern analysis (MVPA) and inter-subject correlations, which do not allow generalization to new stimuli. Further, all past studies that have probed linguistic representations across languages have used at most three languages (see SI S1 for additional discussion of relevant prior work). As a result, past work has left open the question of whether there exists a component in the brain representations of linguistic input that is shared across diverse languages—a question we focus on here.

Leveraging existing fMRI data from a passage listening task in 12 languages (Malik-Moraleda, Ayyash, et al., 2022), we trained encoding models to predict brain responses based on multiple languages and transferred them zero-shot to a new language on which they had not been trained (Study I). Next, we tested the generalizability of the encoding models under more diverse and stringent conditions (Study II): we trained the models on existing fMRI datasets varying in the number of languages, presentation modalities, and types of language stimuli (Blank et al., 2014; Malik-Moraleda, Ayyash, et al., 2022; Pereira et al., 2018; Tuckute et al., 2024a), and then applied them directly to newly collected fMRI data in 9 additional languages, none of which were included in the training data. To foreshadow our key finding, the two studies converged in demonstrating that brain encoding models can robustly predict brain activity across diverse languages, providing evidence for a shared component of linguistic representations. Lastly, we asked what specific properties of language support this cross-lingual generalization. In Studies III and IV, we addressed this question using two complementary approaches—targeted perturbations of the input stimuli and selective removal of linguistic information from the models’ representations—which converged in showing that cross-lingual transfer is primarily driven by meaning, with syntax (and, more generally, form-related properties) playing a more limited role.

## Results

### Study I

Malik-Moraleda, Ayyash, et al. (2022) used fMRI to record brain responses in speakers of diverse languages listening to a ∼4.5-minute-long passage from the book *Alice in Wonderland* (Carroll, 1865) in their native language. We used data from 12 languages (Afrikaans, Dutch, Farsi, French, Lithuanian, Marathi, Norwegian, Romanian, Spanish, Tamil, Turkish, Vietnamese) belonging to 4 language families (Indo-European, Dravidian, Turkic, Austroasiatic). Our sample encompassed high-, mid-, and low-resource languages, differing in the extent to which they have been previously studied in language neuroscience (SI S16). In most languages, participants (n=2 per language) listened to the same passage, but in Afrikaans, Dutch, and French, a different passage from the same book was used. These data were selected from a larger dataset, restricting to languages for which inter-participant blood-oxygen-level-dependent (BOLD) signal correlations in the left-hemisphere language areas (functionally defined; Fedorenko et al., 2010) were reliable (see Methods, “Study I – Time series extraction”).

Our general procedure was similar to that in past studies (e.g., Antonello & Huth, 2024; Aw & Toneva, 2023; Schrimpf et al., 2021; Tuckute et al., 2024a) (**Figure 1A**) and consisted of extracting contextualized text embeddings from the language models for the stimuli in each language, and linearly mapping these representations onto fMRI activity in response to those linguistic stimuli via ridge regression. For each language, we extracted BOLD time series during the passage in the five core areas of the left-hemisphere fronto-temporal language network of each participant (Methods, “fMRI data modeling and definition of functional regions of interest (fROIs)”). To assess the spatial specificity of these effects, we additionally extracted time series from two comparison sets of brain regions: the homotopic right-hemisphere language areas and areas of the multiple-demand (MD) network (Duncan et al., 2020), which plays a limited role in language processing (Fedorenko & Shain, 2021). We averaged the time series across the voxels within each functional region of interest (fROI), yielding a univariate time series per region per participant. We fit our encoding models at the fROI rather than voxel level because our approach requires generalization across individuals and languages, and establishing voxel-wise functional correspondence across individual brains is not trivial given the inter-individual variability in functional topographies (Fedorenko et al., 2010; Nieto-Castañón & Fedorenko, 2012; Fedorenko & Blank, 2020). This fROI-level analysis can establish whether the aggregate functional tuning of each region is shared across languages, but leaves open the question of whether the spatial organization of the tuning of individual voxels within each fROI is similar (see Discussion).

**Figure 1.**
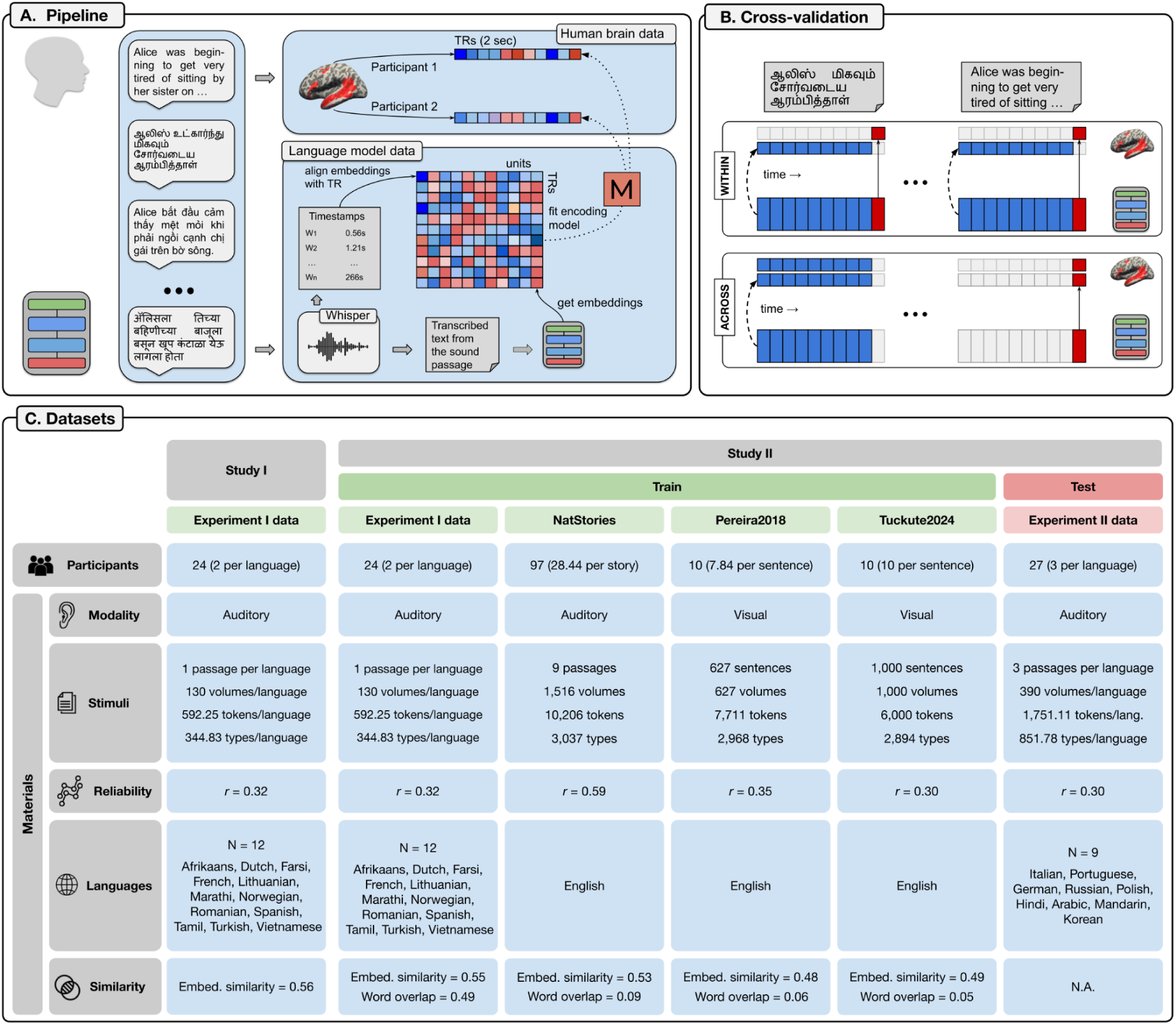
Graphical summary of the study methods. **A. Experimental pipeline:** Passages from the book *Alice in Wonderland* are presented to human participants while in the fMRI scanner (top) and to MNNLMs (bottom). *Human brain data (top):* Two participants listen to the passage in their native language. fMRI time series are obtained from each participant by averaging the responses across voxels in the language areas. *Language model data (bottom):* Written transcriptions of the passages and relative timestamps from the raw audio file are retrieved using the speech-to-text model Whisper-timestamped (Radford et al., 2023; Louradour, 2023). The MNNLM produces embedding representations for the words in the passages, which are then temporally aligned with the fMRI time series. An encoding model is fit to predict fMRI responses on the basis of the MNNLM embeddings. **B. Encoding model cross-validation:** The encoding models are evaluated either on a left-out data subset within each language, or a left-out language. *Within-language cross-validation (top):* The encoding models are trained within each language with 10-fold cross-validation. During training, MNNLM embeddings (large rectangles on the bottom) are mapped onto brain responses (squares on top) in nine data folds using brain data from one of the two participants (blue blocks, dashed arrows). The encoding models then predict brain activity in the remaining data fold using brain responses from the other participant (red blocks, solid arrow). *Across-languages cross-validation (bottom):* The encoding models are trained with cross-validation on nine folds of the brain data in all but one language, and are transferred zero-shot to the held-out fold in the held-out language. **C. Datasets:** Datasets employed in the two studies. For each data source, the table reports the number of participants, presentation modality, information on the stimuli (number of passages/sentences, brain measurements (volumes), tokens, and word types, i.e., unique tokens), split-half reliability, and language coverage. The last row indicates the similarity in the linguistic materials between train and test splits. In the case of Study I, this measure reports the similarity between the word embeddings of each passage across languages. In the case of Study II, it indicates the similarity between the source materials used as training data for the encoding models and the test data (in English), considering both embedding similarity and word overlap. The similarity scores based on the embeddings are averaged across words, sentences/passages, languages, layers, and models.

In addition to fROI-level responses, we also examined responses averaged across the five left-hemisphere language fROIs, and analogously, across their right-hemisphere homotopes and the MD fROIs, because of substantial evidence of functional similarity (Fedorenko et al., 2024a; Assem et al., 2020) and functional connectivity (Blank, Kanwisher, & Fedorenko, 2014; Braga et al., 2020; Shain & Fedorenko, 2025) among the areas within each of these networks. To obtain MNNLM embeddings for the same stimuli (the passages in the 12 languages), we first derived textual transcriptions of the auditory recordings with Whisper-timestamped (Radford et al., 2023; Louradour, 2023), a multilingual speech recognition model that outputs word-by-word timestamps. We then extracted embedding representations of each word in the passage transcriptions from 20 candidate MNNLMs, including models trained on translation (NLLB), contrastive cross-lingual alignment (XLM-Align, InfoXLM), span corruption (i.e., predicting consecutive chunks of text based on bidirectional context; mT5), masked language modeling (mBERT, XLM-R), and causal language modeling (mGPT, XGLM). The models were based on different architectures, encompassing encoder-only, decoder-only, and encoder-decoder models (see Methods, “Language models” and Table 1 for a summary of all the models considered in the study). Next, we aligned the MNNLM contextualized word embeddings with the fMRI time series using the timestamps from Whisper-timestamped and controlling for the hemodynamic delay (**Figure 1A**, bottom). And finally, we fit several linear encoding models to predict the brain responses from the MNNLM embeddings. The encoding models were then evaluated across participants either on left-out data subsets within a language or on left-out languages.

**Table 1.**
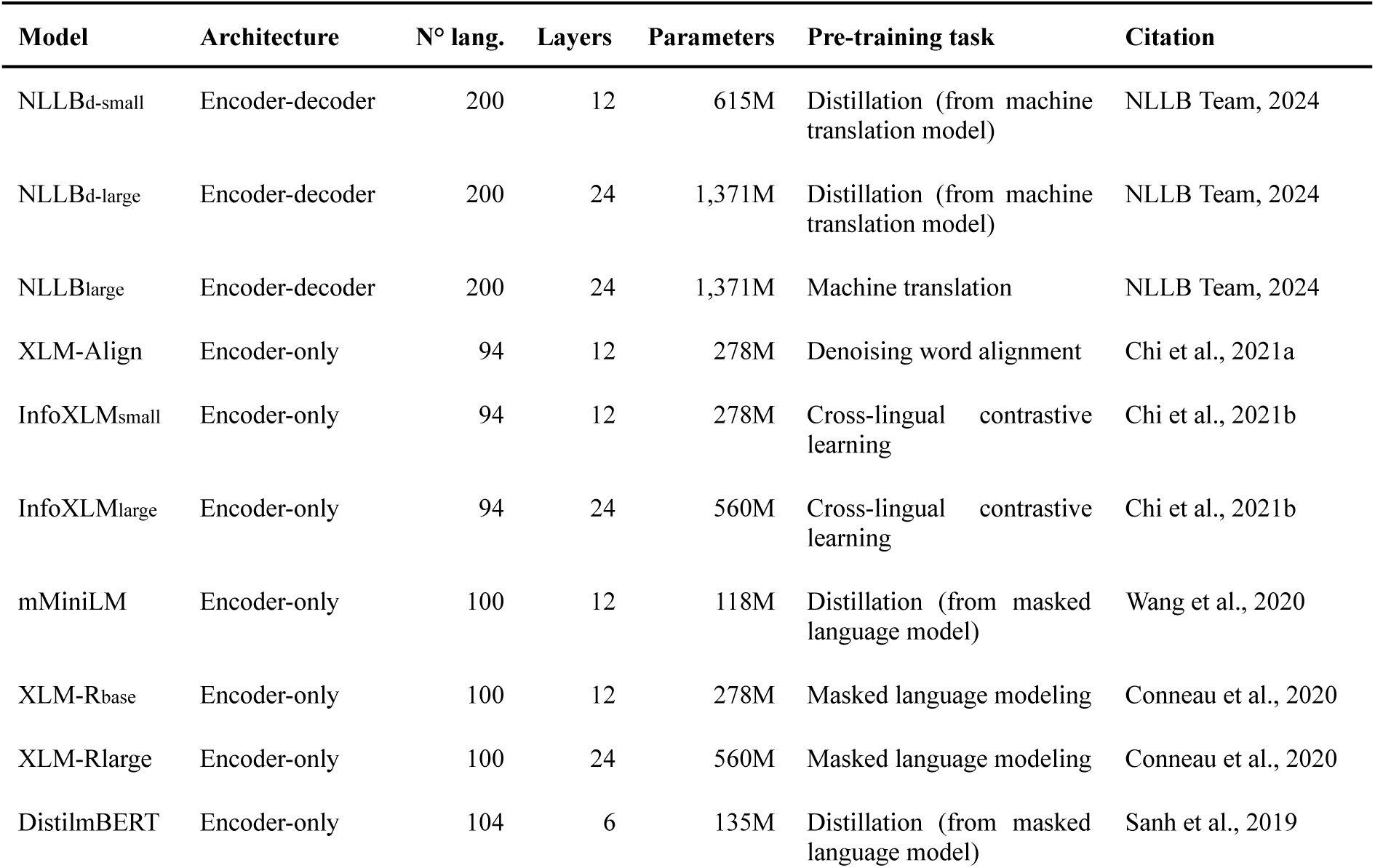
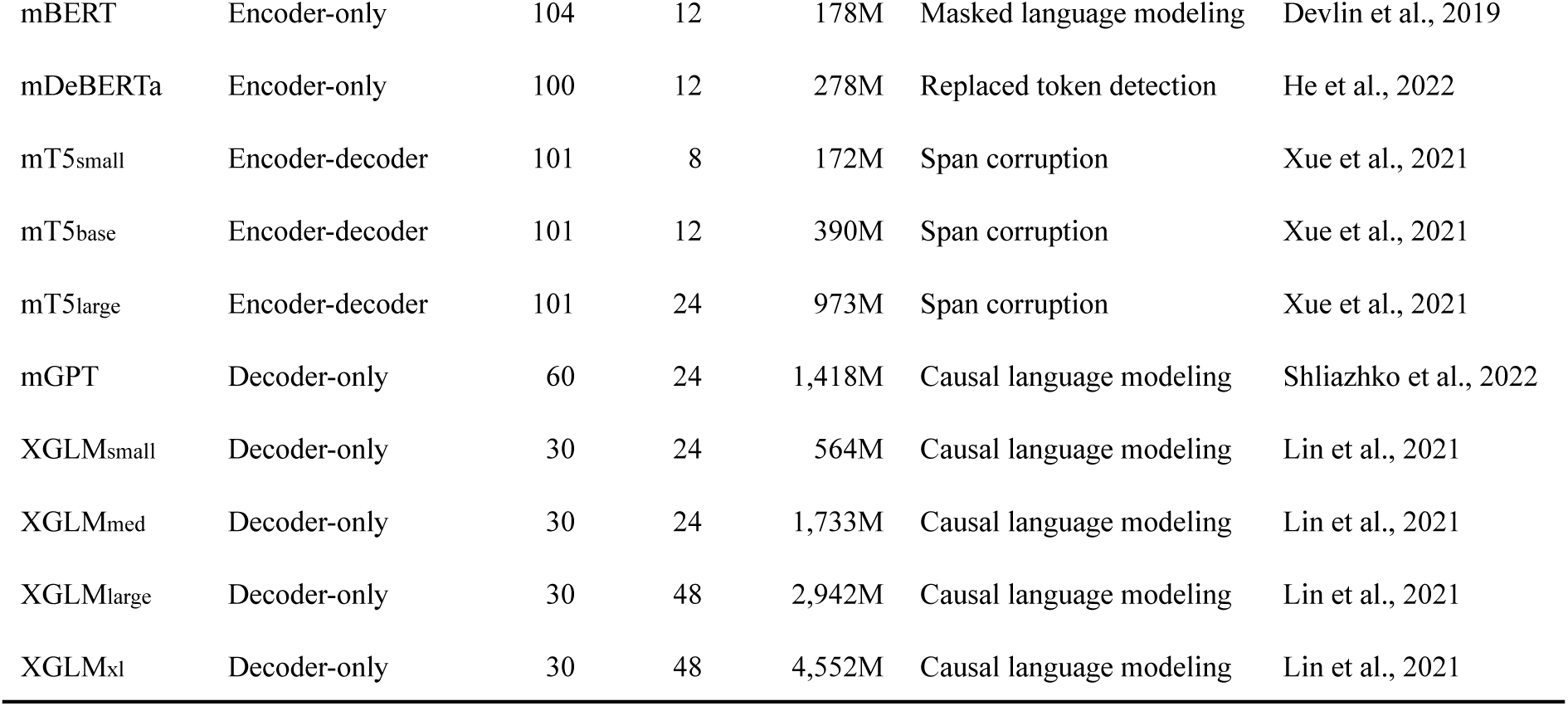
Summary of the MNNLMs employed in the study.

#### Language models predict brain responses in diverse languages

Most past work relating model representations to brain data has relied on data from English and a small number of other high-resource languages, such as Dutch, French, Japanese, and Chinese (Caucheteux & King, 2022; Millet et al., 2022; Nakagi et al., 2024; Sun et al., 2023; Yamashita et al., 2023). To test the ability of language models to capture brain responses for a wider array of languages, we fit an encoding model within each language and evaluated it with a cross-participant design (henceforth the WITHIN condition; see **Figure 1B**, top for a schematic representation of the cross-validation scheme). Encoding models were trained with cross-validation on 90% of the data from one participant and tested on the left-out 10% from the other participant, and vice versa, with results averaged across the two splits. The encoding models’ performance was evaluated as the Pearson correlation between the predicted language network response and the actual fMRI time series. Analyses were performed at two granularities: (i) network-averaged time series—computed by averaging the five left-hemisphere language fROIs (and, analogously, the five right-hemisphere homotopes and the 20 MD fROIs)—and (ii) individual fROI time series analyzed separately. We calculated statistical significance by comparing the performance of the encoding models with a circular shift baseline in which the fMRI time series was shifted by a variable number of TRs, thereby preserving the autocorrelation structure while disrupting alignment with the stimulus (Methods, “Significance testing”).

Our results show that MNNLMs can successfully predict the language network’s response to diverse languages, as recorded with fMRI (**Figure 2A**, top; see SI S2 for the results in each individual language), significantly outperforming the circular shift baselines for nearly all models (across-model average *r* = 0.23, *SE* = 0.01, all *p* < 0.05 apart from NLLBd-small).

**Figure 2.**
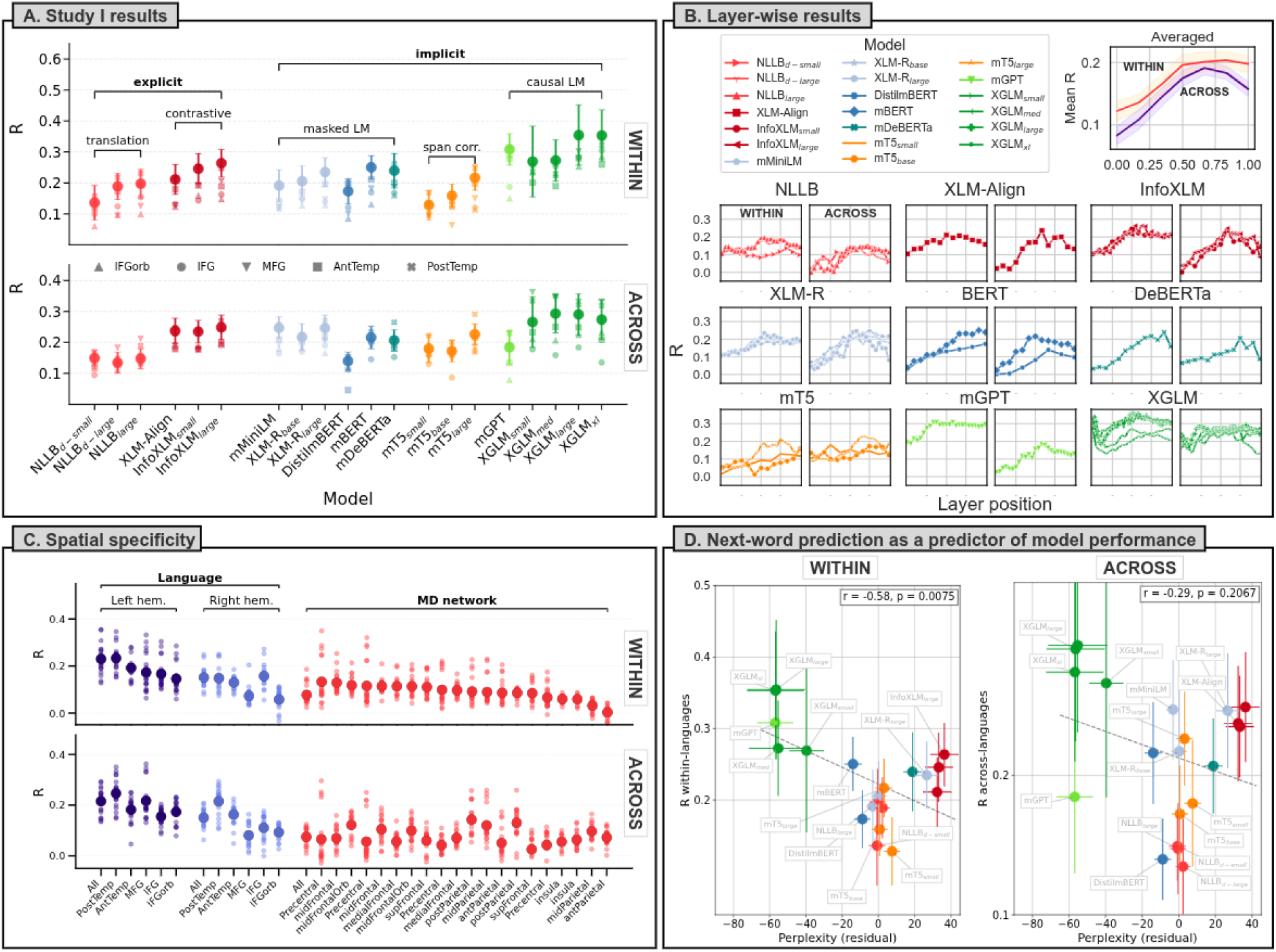
Summary of the results of Study I. **A. Study I results:** *Top:* Best-layer WITHIN encoding performance by model, averaged across languages. The models are first grouped based on whether they are exposed to implicit or explicit cross-lingual signal during training, and then by their training objective. Models in the same colors belong to the same model family. The larger dots indicate the encoding performance in the language network, whereas the smaller markers indicate the specific fROIs. The error bars indicate the standard error of the encoding performance across languages. Causal (or auto-regressive) language models achieve the strongest brain encoding performance; within each model family, larger-capacity models are more effective in predicting brain activity. *Bottom:* Best-layer ACROSS encoding performance by model type. See SI S5 for the results obtained with the median-performing layer in the two conditions. **B. Layer-wise results:** Encoding performance in the language network by layer depth. The subplot on the top right reports the layer-wise encoding scores averaged across models; the shaded areas around the lines indicate the standard error of encoding performance across models. The layer positions (*x*-axis) are normalized (i.e., divided by the total number of layers in the models). The remaining subplots indicate the encoding performance by layer for all models. For each model family (NLLB, XLM-Align, etc.) the first plot reports the layer-wise results obtained in the WITHIN condition, whereas the second plot depicts the scores obtained in the ACROSS condition. Encoding scores are averaged across languages. **C. Spatial specificity:** Encoding performance by fROI in three networks of interest: left-hemispheric areas of the language network (dark blue), right-hemispheric language areas (light blue), and areas of the Multiple Demand network (red). Both in the WITHIN (top) and ACROSS conditions (bottom), encoding performance is higher in the left-hemisphere core language regions than in either set of control areas. **D. Next-word prediction as a predictor of model performance:** Relationship between the models’ next-word prediction abilities and encoding performance, both in the WITHIN (left) and ACROSS (right) conditions. Next-word prediction (*x*-axis) is operationalized as perplexity on held-out data subsets, averaged across languages. Lower perplexity values indicate better next-word prediction accuracy. Encoding performance (*y*-axis) is measured as the best-layer encoding performance for each model considered. The error bars indicate the language-wise standard error of the mean on the two axes. The gray dashed line indicates the best linear fit. Note that the *y-*axes use different scales.

At the network level, causal (or auto-regressive) MNNLMs – i.e., models trained to predict the next word in a text sequence – achieve the best brain encoding performance, generally outperforming models trained on translation, cross-lingual contrastive learning, span corruption, or masked language modeling, in line with previous observations based on data from English (Schrimpf et al., 2021). Furthermore, within each model family, larger-capacity models outperform their smaller counterparts. Indeed, the best average performance is obtained by XGLMxl (*r =* 0.35, *SE =* 0.08) and XGLMlarge (*r =* 0.35, *SE =* 0.10) followed closely by other large-scale auto-regressive models (mGPT, *r =* 0.31, *SE =* 0.05; XGLMmed, *r =* 0.27, *SE* = 0.07). Encoder-decoder models, trained on translation (NLLB family) or span corruption (mT5 family), are associated with lower average performance, demonstrating reduced effectiveness in capturing language-specific brain responses compared to other model types.

Analyses of individual language fROIs revealed patterns that were broadly consistent with the network-level pattern and with each other (**Figure 2A**, small markers; see also **Figure 2C**), with reliable encoding in each of the five left-hemisphere regions. Encoding performance is slightly stronger in the temporal compared to the frontal regions, plausibly because of better signal in those areas (see Figure 4 for the split-half correlation in the time series for each fROI). For example, the average encoding score in the posterior temporal language fROI reaches *r =* 0.23 across models (ranging from 0.15 to 0.35, *SE =* 0.03–0.07), whereas encoding in the inferior frontal language fROI only reaches *r* = 0.17 across models (range: 0.10–0.30, *SE =* 0.04–0.09) (see SI S8 for details on each fROI). These fROI-level results show that the network-level effects are not driven by a single fROI, but instead reflect effects that are present across the fronto-temporal language network. By comparison, the right-hemisphere homotopic regions show substantially weaker effects (across-model mean *r =* 0.15, *SE =* 0.009, compared to *r* = 0.23, *SE* = 0.01 in the left hemisphere), and the MD network effects are even lower (mean *r =* 0.08, *SE =* 0.009). These findings establish that language information is not ubiquitously present across the brain and mirror past work on English (e.g., Tuckute et al., 2024a).

The WITHIN encoding performance is critically influenced by the depth of the layer within the model. **Figure 2B** displays the encoding performance scores for each layer, normalized by their position within the model. For most model families, intermediate layers (as seen in XLM-Align, InfoXLM, XLM-R, DeBERTa) and deeper layers (as observed in mBERT, mT5, mGPT) demonstrate the highest predictivity in relation to the fMRI signal, also in line with prior work (Schrimpf et al., 2021; Antonello et al., 2023). The layer-wise performance patterns are less clear for the NLLB and XGLM models. Encoding scores averaged across models increase until a normalized depth of ∼.70, where they reach a plateau (**Figure 2B**, top right).

#### Encoding models can be transferred across languages

To test for the existence of a shared component of linguistic representations *across* languages, we trained the encoding models on the data in all but one language, and tested them zero-shot on the left-out language (the ACROSS condition; see **Figure 1B**, bottom for a visualization of the cross-validation scheme; see SI S14 for a more stringent test where encoding models are trained in a single language and tested in each of the remaining ones). Similar to the WITHIN analyses, models were fit and evaluated in a cross-participant design.

Encoding models can be successfully transferred zero-shot to unseen languages, highlighting the strong generalizability of the link between distributed language representations learned from statistical word co-occurrence patterns and brain responses in the human language network (**Figure 2A**, bottom). At the network level (averaging across the five left-hemisphere language fROIs), the across-model mean is *r =* 0.22, *SE =* 0.011, well above the circular-shift baseline (all *p* < 0.001). Large auto-regressive models obtain the strongest performance (XGLMmed: *r =* 0.29, *SE =* 0.06; XGLMlarge: *r =* 0.29, *SE =* 0.07; XGLMxl: *r =* 0.27, *SE =* 0.06; XGLMsmall: *r* = 0.27, SE = 0.08), while encoder-decoder models obtain lower transfer performance, confirming their limited capacity in capturing brain responses compared to other model classes (as already shown in the WITHIN condition).

Analyses of individual left-hemisphere language fROIs again mirror the pattern obtained for the network as a whole, showing consistent transfer (**Figure 2A**, small markers; **Figure 2C**). Effects are again strongest in the posterior temporal fROI (average *r =* 0.25 across models; range: 0.15–0.35; *SE* = 0.03–0.07), and reliable but slightly weaker in the other fROIs (see SI S8 for details). By comparison, the right-hemisphere homotopic regions showed substantially weaker effects (across-model mean *r =* 0.15, *SE =* 0.01, compared to *r* = 0.22, *SE* = 0.01 in the left hemisphere), and the MD network lower still (mean *r =* 0.08, *SE =* 0.01).

Layer depth within the model remains a significant factor in the ACROSS condition (**Figure 2B**). Intermediate-to-deep layers show higher levels of predictivity, paralleling the trends observed in the WITHIN encoding models and prior work. Encoding scores averaged across models display an inverse U-shaped relation with layer depth, increasing until a normalized depth of ∼.70 followed by a decrease (**Figure 2B**, top right).

#### Next-word prediction ability explains inter-model differences in within-language encoding performance

Previous research in computational language neuroscience has shown that language models with better next-word prediction abilities provide a better basis for brain encoding models (Schrimpf et al., 2021; Caucheteux & King, 2022; Hosseini et al., 2024). To test whether this observation extends to diverse languages, we evaluated the next-word prediction abilities of our 20 models by training a ridge classifier on top of the MNNLM embeddings to predict the next word in a multilingual corpus from Wikipedia. Next-word prediction scores for each model were quantified as the classifiers’ cross-validated perplexity, averaged across languages (lower perplexity corresponds to better next word prediction ability). Extending prior findings from English (Schrimpf et al., 2021; Hosseini et al., 2024) and Dutch (Caucheteux & King, 2022) to a wider array of languages, next-word prediction ability showed a significant relationship with encoding performance in the WITHIN condition (*r* = -0.58, *p* < 0.01; **Figure 2D**), indicating that more accurate language models provide a stronger basis for neural encoding. This pattern did not hold for the ACROSS condition (*r* = -0.29, n.s.), but the difference between the two correlations was not significant. The correlation in the WITHIN condition should not be over-interpreted as evidence that the brain’s language network is optimized for next-word prediction. As argued by Antonello & Huth (2024), next-word prediction produces broadly generalizable representations, and this property of the representations may be the critical mediating factor (see also Tuckute et al., 2024b for discussion).

### Study II

In our first study, we found that the relationship between language representations in artificial and biological systems is sufficiently general to be successfully transferable across languages. In our second study, we devised an even stronger test for the cross-lingual transferability of the encoding models. We fit the encoding models employing fMRI data from Study I (passage listening in 12 languages) and from three additional fMRI datasets, all using English stimuli presented via auditory or visual modalities. Then, we collected new fMRI data on 9 additional languages (non-overlapping with those in Study I), using the same paradigm of listening to passages from *Alice in Wonderland*, and then directly applied the trained models zero-shot to these newly collected brain responses. The three additional training datasets included two datasets where participants read English sentences (*Pereira2018*, Pereira et al., 2018, Experiment 2 and 3; *Tuckute2024*, Tuckute et al., 2024a) and an fMRI dataset where participants listened to naturalistic stories in English (*NatStories*, expanded from Blank et al., 2014; parts of this datasets were used in Shain, Blank et al., 2020, Shain et al., 2022, and Paunov et al., 2022; see Methods, “NatStories” for details). The datasets thus varied in the languages covered (one vs. multiple languages), type of stimuli employed (sentences vs. passages/stories), and presentation modality (listening vs. reading). **Figure 1C** summarizes the information about the datasets, including details on the participants, stimuli, response reliability, and similarity of the language materials to the test set. In contrast to Study I, where all participants listened to passages from the same book, in Study II, the source materials on which we trained the encoding models involve diverse language stimuli with little word overlap with the test materials and lower similarity in the embedding representations.

We collected new fMRI data from native speakers of 9 languages (Italian, Portuguese, German, Russian, Polish, Hindi, Arabic, Mandarin, Korean) belonging to 4 language families (Indo-European, Afroasiatic, Sino-Tibetan, Koreanic). For each language, three native speakers listened to three passages (∼4.5 min each). As in Study I, the participants’ language areas were identified with an independent language localizer task (Fedorenko et al., 2010; Malik-Moraleda, Ayyash et al., 2022), and we extracted the responses during the critical passage listening task from these functionally defined areas and averaged across them, and then averaged the time series across participants (two or three, depending on the reliability of the time series; see Methods, “Study II – Time series extraction”). Similarly to Study I, in Study II we averaged responses across voxels within each fROI and then across fROIs (although we report in SI S8 the results for the individual fROIs in addition to the network-level results). Differently from Study I, we additionally averaged across participants, resulting in a single response per sentence (*Pereira2018*, *Tuckute2024*) or a single time series per passage/story (*Study I data*, *NatStories*). Our general approach was identical to Study I with respect to pre-processing, extraction of time series, temporal alignment of MNNLM embeddings and neural data, and model fitting, with the only difference that for the sentence-level datasets (*Pereira2018, Tuckute2024*), which involved no temporal structure, statistical significance was evaluated against a shuffled baseline instead of a circular shift baseline.

#### Encoding models can be transferred to new languages across types of language materials and presentation modalities

The results of Study II show that encoding models trained on one language or a set of languages can be successfully applied to new languages even across variation in type and content of stimuli and presentation modalities (**Figure 3A**). The strongest transfer performance is obtained by encoding models trained to predict brain responses in participants listening to passages or stories (Study I and *NatStories*; **Figure 3A**, top). All 20 encoding models based on the data from Study I significantly outperform the randomized baseline, reaching an encoding performance as high as *r* = 0.17 (*SE* = 0.06) in the case of XLMRlarge. The encoding models trained on the *NatStories* data achieve an even stronger generalization performance, as all of them yield encoding scores significantly above chance, with correlations of up to *r* = 0.21 (*SE* = 0.10) with the actual brain responses for the mGPT model. We attribute the strong performance of the encoding models trained on *NatStories* to the high reliability of the fMRI responses in this dataset, which is likely due to a relatively large number of participants per story (**Figure 1C**). Our findings also reveal that encoding models trained on data from sentence-reading tasks (*Tuckute2024, Pereira2018*) can predict brain responses to passage-listening tasks (**Figure 3A**, bottom). This cross-modality/cross-stimulus type of cross-lingual transfer highlights the robustness of the encoding models and suggests a deep alignment between the responses evoked by reading and listening in the language network, in line with a large body of prior work (e.g., Deniz et al., 2019; Fedorenko et al., 2010; Regev et al., 2013; Vagharchakian et al., 2012). In particular, models trained on the *Tuckute2024* dataset yield strong cross-modality and cross-lingual performance when mapped onto listening-based fMRI data, with all tested models exhibiting statistically significant predictive performance. The highest encoding score of the models trained on this dataset is achieved by mGPT, with a correlation of *r* = 0.17 (*SE* = 0.06) between predicted and observed brain responses. Models trained on the *Pereira2018* dataset exhibit a more modest cross-modality transfer performance, with 15 of the 20 models predicting brain responses above chance, and the best-performing model, mGPT, achieving an encoding score of *r* = 0.11 (*SE* = 0.03). The zero-shot generalization ability of MNNLM-based encoding models—shown in Study I and replicated and extended in Study II—indicates that MNNLMs capture properties of language processing that remain consistent across a range of linguistic and experimental contexts. This transferability highlights the potential of brain encoding models to reveal general principles of brain function related to language.

**Figure 3.**
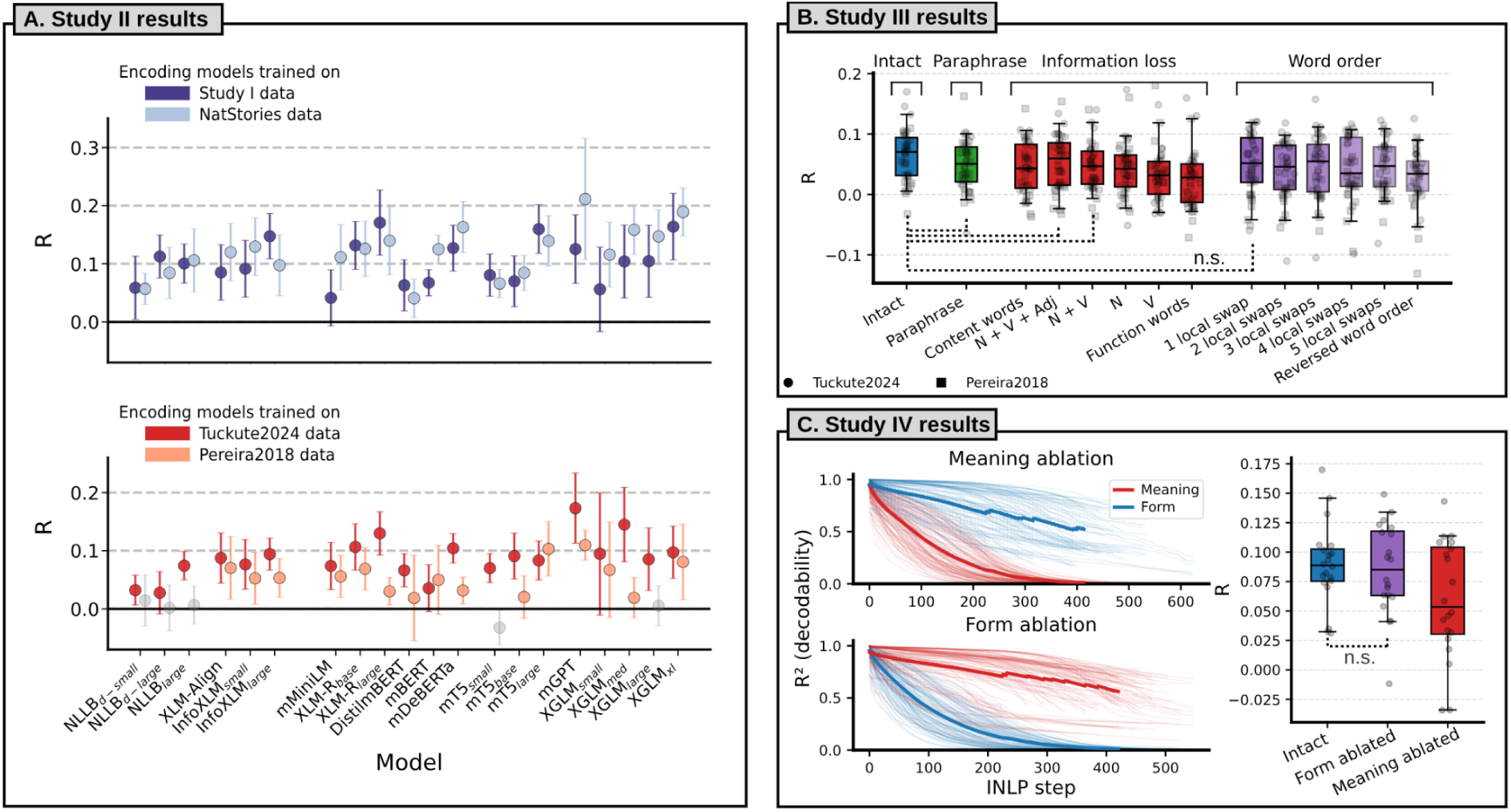
Summary of the results of Studies II, III, and IV. **A:** Results of Study II. The dots indicate how well the models trained on the independent data sources (Study I, *NatStories*, *Pereira2018*, *Tuckute2024*) transferred to the new data we collected. The encoding scores are averaged across passages and languages (see SI S3 for the results in the individual languages); the error bars indicate the standard error of the encoding performance across languages. The models are grouped according to the taxonomy defined for Study I. The semi-transparent dots indicate models whose encoding performance did not reach statistical significance. Top: Transfer performance of the encoding models trained to predict brain responses to passages/stories presented auditorily (employing the data from Study I and *NatStories*). Bottom: Transfer performance of the encoding models trained to predict brain responses to sentences presented visually (employing the data from the *Tuckute2024* and *Pereira2018* datasets). **B:** Results of Study III. Performance of the encoding models trained on the “perturbed” versions of the *Tuckute2024* and *Pereira2018* datasets in transferring to the new data we collected. The results within each perturbation type are averaged across models and across the two datasets. The individual data points indicate the results obtained by each model in each dataset. All “perturbed” encoding models performed significantly worse than the intact encoding model apart from those marked with the dotted lines (*n.s.*; N + V + Adj, N + V, Paraphrase, 1 local swap). **C:** Results of Study IV. The two line plots on the left indicate the linear decodability of the seven meaning-related features (red) and form-related features (blue) (*y* axis) at different steps of the Iterative Null-space Projection method (INLP; *x* axis), both for meaning ablation (top) and form ablation (bottom). The thin lines indicate decodability of individual model × feature combinations, whereas the thick lines indicate grand averages. Meaning ablation makes meaning-related features undecodable while comparatively preserving form-level information (top); conversely, form ablation makes form-related features undecodable while comparatively preserving meaning-level information (bottom). The boxplot on the right reports the performance obtained by the MNNLMs (gray dots) trained on the *Tuckute2024* dataset in transferring to the new data collected for Study II. Encoding models based on form-ablated embeddings performed comparably to those trained on intact embeddings, whereas meaning-ablated embeddings significantly reduced transfer performance.

### Study III

Through studies I–II, we have shown that encoding models trained in one language transfer to other languages, which implies the existence of a shared cross-lingual component in the responses of the language network. What is the nature of this shared component? First, we asked whether the shared component may have to do with low-level stimulus features. In particular, we examined how well brain responses to language could be predicted by a simple baseline of word length and word frequency—features that explain substantial variance in linguistic processing difficulty as reflected in reading times (Clifton et al., 2016; Kuperman, Schroeder, & Gnetov, 2024), and features such as word rate and word onset. Encoding models based on this baseline achieved low predictivity, demonstrating that MNNLM encoding models capture neurally relevant information beyond these low-level features (SI S6; see also de Varda et al., in prep.).

Next, to narrow in within the space of higher-level linguistic features, we applied targeted perturbations to the stimuli and measured the effect on the models’ generalization (following the approach of Kauf, Tuckute, et al., 2024). Perturbations came in three flavors—a paraphrase (produced by a large language model, Llama 4 Maverick), removal of certain word categories (“information loss” manipulations), and word order changes—and varied in the degree to which they affected the representation of the sentence meaning. In particular, a paraphrase, removal of only the function words, and minor word order changes via local swaps largely preserve the meaning of the sentence, but removal of all content words or a complete reversal of the word order substantially affect the compositional meaning. Encoding models were trained on embeddings of perturbed English sentences from the *Pereira2018* and *Tuckute2024* datasets, and tested for generalization in the nine other languages, using the same procedures and data as Study II.

As expected, cross-lingual predictivity was highest for intact sentences (*r =* 0.065, *SE =* 0.007). Paraphrases yielded similar performance (*r =* 0.049, *SE =* 0.007, 74.36% of intact). Performance was also not too strongly affected by information loss manipulations that preserved most of the lexical-semantic content (all content words preserved: *r =* 0.045, *SE =* 0.007, 69.54%; nouns, verbs and adjectives preserved: *r =* 0.053, *SE =* 0.007, 81.72%), or by word order manipulations that involve local word swaps (with *r* ranging from 0.043 to 0.049, *SE ≈* 0.008, 66.21–75.44%). However, performance showed a substantial drop when only nouns were preserved (*r =* 0.042, *SE =* 0.007, 64.41%) or only verbs (*r =* 0.034, *SE =* 0.007, 52.35%). And performance was lowest when only function words were preserved (*r =* 0.025, *SE =* 0.007, 37.61%) or when the word order was fully reversed (*r =* 0.028, *SE =* 0.008, 42.72%). These results show that cross-lingual transfer is primarily driven by compositional semantic content: when too many content words are removed or when the order of the words is changed so much that the structure of the sentence is difficult to reconstruct, transfer performance drops to very low levels.

### Study IV

The perturbation analyses in Study III demonstrated that cross-lingual transfer is primarily driven by compositional semantic content. However, the perturbation approach has a potential limitation: because the MNNLMs were trained on intact sentences, perturbed inputs are increasingly out-of-distribution, and the resulting embeddings may be degraded in ways that are not specific to the removal of particular linguistic information. As a result, we conducted a complementary analysis that operated directly on the embedding representations, instead of on the stimuli.

We implemented a variant of the Iterative Null-space Projection method (INLP; Ravfogel et al., 2020), a procedure that iteratively identifies linear subspaces that encode a target set of features and removes those subspaces from the representations. We defined two sets of seven features each to be removed from the MNNLM embeddings: a set of meaning-related features (human sentence-level ratings of imageability, valence, arousal, real-world plausibility, and the presence of three kinds of content: others’ thoughts, physical interaction, and places; data from Tuckute et al., 2024a) and a set of form-related features (PCFG surprisal, mean dependency length, mean word frequency, and mean word length, and human ratings of grammaticality, frequency, and conversational frequency). For each of the 20 MNNLMs, we extracted sentence-level embeddings for the *Tuckute2024* stimuli at the best-performing layer identified in Study I and used in Studies II and III, and then applied INLP to produce two residualized versions of the embeddings: one from which meaning-related information is removed, and one from which form-related information is removed. We confirmed that INLP successfully reduced the linear decodability (R^2^) of the targeted features to below 0.01 for all models, while comparatively preserving the decodability of the non-targeted features (**Figure 3C**, left).

We then trained encoding models on the intact, meaning-ablated, and form-ablated embeddings and evaluated their ability to transfer zero-shot to the 9 languages from Study II. Removing meaning-related information from the embeddings led to a substantial reduction in cross-lingual transfer performance, with meaning-ablated models conserving only 67.0% of the intact encoding performance (meaning-ablated: *r* = 0.060, *SE* = 0.050; intact: *r* = 0.089, *SE* = 0.035; *z* = −3.24, *p* = 0.001). In contrast, removing form-related information had a negligible effect on transfer, with form-ablated models conserving 95.5% of the intact performance (form-ablated: *r* = 0.085, *SE* = 0.039; *z* = −0.46, n.s.). The reduction in transfer performance was significantly greater for meaning ablation than for form ablation (*z* = −2.77, *p* = 0.006). This is in contrast with the results obtained in English with within-language cross-validation, where meaning and form ablation both showed substantial impact on performance (SI S17). Together, Studies III and IV provide complementary converging evidence (from targeted perturbations applied to the stimuli or to the associated embeddings directly) that the shared component of brain responses across languages is primarily related to meaning.

## Discussion

At the core of the language sciences lies a fundamental tension between the extensive variety of human languages and the uniformity of the biological tissue that supports their use. Indeed, the human language network—an interconnected set of left-lateralized frontal and temporal brain regions—supports the processing of all human languages tested so far, irrespective of their profound surface differences that span most levels of linguistic analysis (Chee et al., 1999; Illes et al., 1999; Rueckl et al., 2015; Malik-Moraleda, Ayyash, et al., 2022), and even including constructed languages, such as Klingon (Malik-Moraleda et al., 2025). This common neural substrate for language processing presumably emerged through biological and cultural evolution in response to similar functional demands and constraints. But beyond broadly cross-lingually similar anatomical and functional properties of the language network, are linguistic representations similar across languages? Recent technological advances in the form of multilingual neural network language models (MNNLMs) have made it possible to derive quantitatively specified linguistic representations in a language-general format. Building on these advances, we first replicated the findings from English and a handful of other high-resource languages (e.g., Schrimpf et al., 2021; Caucheteux & King, 2022; Millet et al., 2022; Nakagi et al., 2024; Sun et al., 2023; Yamashita et al., 2023) showing that encoding models based on these MNNLMs capture neural responses to language in 21 diverse languages (12 in Study I and 9 in Study II; see SI S4). Next and critically, we tested whether the link between language representations and brain responses (i.e., our encoding models) can be transferred across languages. Our results show that indeed the features encoded by MNNLMs can capture brain responses across languages, indicating the existence of a substantial component of the brain response to language that is governed by language-general principles.

What kind of linguistic information supports the cross-lingual transfer of the encoding models? Contextualized word embeddings are high-dimensional language representations that encode both surface (form-related) features of words and sentences, and meaning features, both lexical and compositional (Jawahar et al., 2019). Our results demonstrate that not all the information encoded in the MNNLMs’ embeddings can be successfully transferred across languages: encoding models perform better in the WITHIN condition than in the ACROSS condition (Study I), despite being trained on substantially less data. Identifying the exact subset of features that support cross-lingual transfer is not trivial and would likely require carefully controlled comparisons across many language pairs. Here, we consider two broad possibilities—whether the transfer effects are primarily driven by linguistic form versus by linguistic meaning.

Linguistic form is a compelling candidate for mediating cross-lingual transfer because human languages share a number of structural regularities driven by general cognitive and communicative pressures (see Fedorenko et al., 2024b; Gibson et al., 2019; Kemp, Xu, & Regier, 2018; Levshina & Moran, 2021 for reviews). For example, in all languages, frequent and predictable words tend to be shorter (Zipf, 1949; Piantadosi et al., 2011) and syntactic dependencies tend to be local (Futrell, Mahowald, & Gibson, 2015; Gildea & Temperley, 2010). Critically, brain activity in the language network is strongly sensitive to these lexical and syntactic features (e.g., Heilbron et al., 2022; Lopopolo et al., 2021; Sánchez, Carreiras, & Paz-Alonso, 2023; Shain, Blank et al., 2020; Shain et al., 2022; Tuckute et al., 2024a). By encoding these statistical regularities, MNNLMs might therefore capture form-based features that modulate brain responses in a similar way across languages. In one analysis, we ruled out the possibility that the simplest form-based features (word length and frequency) and speech features, such as word rate, explain cross-lingual transfer (SI S6). Moreover, we performed a series of analyses aimed at testing the effects of form-based inter-language similarity (including phonetic, phonological, and syntactic similarity) on transfer. The results (reported in SI S14) show no evidence of form-based similarity at any level on transfer. These null results should be interpreted with caution due to potential measurement noise (including variability in data reliability across languages, which we show in SI S14 to be a strong predictor of transfer performance) and limited statistical power in comparing encoding performance between languages (SI S15). Nevertheless, these results provide tentative evidence that form-level features may not be the primary determinant of cross-lingual encoding transfer. This interpretation is further supported by the perturbation analyses in Study III, which showed that function words alone carried little information for transfer and that local word order swaps did not have much of an effect, and by the results of Study IV, where selectively removing form-related information from the MNNLM embeddings had a negligible effect on cross-lingual transfer.

Alternatively, cross-lingual transfer could be supported by higher-level information related to sentence meanings: after all, expressing complex meanings is the core objective of language use, and all human languages are capable of expressing diverse meanings about the outer and inner worlds. Although languages greatly vary in their surface features, cross-lingual variation in semantics, while attested (Everett, 2005; Frank et al., 2008; Majid et al., 2018; Berlin & Kay, 1969; Gibson et al., 2017), is comparatively limited relative to the massive overlap across languages in the concepts they can express (Youn et al., 2016; Liang, Xu, & Ran, 2024). Three aspects of our results point to compositional semantic information playing a critical role in the transfer. First, Study III demonstrated that cross-lingual transfer remains high in conditions that largely preserve the meaning of the sentence: paraphrases and sequences of content words in the right order generalized almost as well as the original sentences. The reverse word-order condition further suggests that the transfer effects go beyond the individual word meanings: the same words presented in an order where the compositional meaning is difficult to infer led to low transfer performance. Second, Study IV provided converging evidence by directly removing meaning- or form-related information from the embedding representations. Removing meaning-related information substantially reduced cross-lingual transfer, whereas removing form-related information had a negligible effect. And third, in the ACROSS condition, intermediate-to-deep model layers yield the best encoding performance. In contrast to early model layers, which encode low-level linguistic features such as morphological structure and parts of speech, deeper layers represent higher-level information such as compositional semantics (Belinkov et al., 2020; Niu et al., 2022; Tenney et al., 2019). These findings converge with previous observations that sentence meanings are the main contributor to the alignment between representations from language models and fMRI recordings (Caucheteux et al., 2021; Gauthier & Levy, 2019; Kauf, Tuckute et al., 2024). Nevertheless, the role of form-based features should be investigated more thoroughly in future work.

If the successful transfer of the encoding models relies on somewhat form-independent semantic representations in the brain, a critical question concerns the degree of abstractness of such representations. In other words, are the shared responses that we documented specific to language? Or would they be similarly well captured by semantic models derived from other modalities, such as vision (Kriegeskorte, 2015)? Because language is often used to talk about the physical world, relations between words tend to reflect relations between the words’ referents (de Varda, Petilli, & Marelli, 2025; Günther et al., 2020; Johns & Jones, 2012; Louwerse, 2011). Indeed, recent work has shown that semantic alignment emerges naturally across modalities, as both language and visual representations encode statistical regularities in the underlying reality (Roads & Love, 2020; Huh et al., 2024). In line with this similarity between linguistic and visual statistics, i) representations from neural network language models can successfully predict brain responses in visual cortical areas (Doerig et al., 2022), even in macaques (Conwell et al., 2024), ii) representations from vision models can predict brain responses in linguistic areas (Saha et al., 2025; Small et al., 2025), and iii) brain encoding models based on multimodal transformers (Xu et al., 2023) can be transferred across language and vision in putative semantic brain areas (Tang et al., 2024). It is important to note, however, that the language areas we are focusing on here are not amodal semantic areas: they respond much more strongly to semantic information expressed in language compared to information carried by meaningful non-linguistic stimuli, such as pictures or videos (Ivanova et al., 2021; Sueoka, Paunov et al., 2024). As a result, more work is needed to understand the nature of semantic representations in brain areas that show a strong preference for a particular representational format, such as the language areas or high-level visual areas, versus in general semantic areas (Lambon Ralph et al., 2016; Ryskina et al., 2025: Ivanova et al., 2025).

Beyond showing cross-lingual transfer effects, our results corroborate previous observations from the computational neuroscience literature using a wider array of languages. Against the backdrop of English-centricity in the current literature, we show that language models can successfully account for brain responses even in low-resource languages, such as Marathi and Lithuanian. These findings resonate with the results of Hosseini et al. (2024), who showed that language models align with brain responses even if trained on developmentally plausible amounts of data. More generally, they reinforce the idea that brain responses to language are modulated by its distributional statistics, which, in turn, are well captured by computational language models. Furthermore, not only can language models account for brain responses in diverse languages, but the model properties that have been shown to affect predictive accuracy for English extend to the languages in our sample. In particular, larger models outperform smaller models in predicting brain activity, underscoring the representational richness of human linguistic representations; and better next-word prediction performance of MNNLMs is associated with stronger ability to predict neural responses. Regarding this last finding, recent work cautions against interpreting it as evidence that the brain itself is engaged in predictive processing: such training may simply be an efficient way to learn broadly generalizable representations (Antonello & Huth, 2024).

The current study suffers from two main limitations. First, our analyses operate at the level of functionally defined regions of interest (fROIs). Although these fROIs are defined in individuals and are much smaller than the anatomical areas often used in neuroscience studies, they still comprise dozens of voxels (between 37 and 294 voxels for the five core language areas) and likely have complex internal structure. Successful cross-lingual transfer at this fROI level implies that the aggregate functional tuning of each region (the relative weighting of different linguistic dimensions in the aggregate response of each fROI) is similar across languages. However, cross-lingual transfer does not require that the spatial arrangement of the voxels’ tuning within each fROI be identical: the same combination of linguistic features may be organized differently across voxels within a fROI in speakers of different languages. Understanding the fine-grained spatial organization of cross-lingual representations will require substantially larger amounts of data per participant per language so as to support stable voxelwise encoding models (as in Huth et al., 2016; LeBel et al., 2021). Further, evaluating the similarity of the fine-grained spatial arrangements across individuals will require some methods to establish meaningful correspondence between voxels across participants whose fROIs differ in spatial configuration.

And second, in spite of being able to identify cross-lingual similarities in brain responses, our design is not equipped to evaluate fine-grained differences in brain encoding across languages (see SI S14 and S15). In particular, the reliability of the fMRI time series varies considerably across languages in our dataset, and this variability is itself a significant predictor of cross-lingual transfer performance (SI S14). Furthermore, the MNNLMs used in this study were not developed to conduct rigorous cross-lingual comparisons, as their pre-training data distribution was highly unbalanced across languages. MNNLMs typically mitigate resource imbalances through strategies such as upsampling data for low-resource languages (Conneau et al., 2020; Devlin et al., 2019; Xue et al., 2021), but high-resource languages remain over-represented in the pre-training corpora, leading to inherent differences in representation quality across languages. Although there have been recent efforts to develop comparable monolingual models in hundreds of languages (Chang et al., 2024), extending a similar standardization to multilingual models has not yet been accomplished, yet is a necessary step for robust, controlled comparisons. Additionally, future studies evaluating inter-language differences will need to adopt a “deep sampling” approach to fMRI data collection, testing large groups of speakers for each language and carefully matching them on potentially relevant factors. This is crucial because cross-lingual differences are likely to be subtle and must exceed the considerable variability among the speakers of the same language to be reliably detected (Malik-Moraleda, Ayyash, et al., 2022).

What could these cross-lingual differences in encoding look like? There are several factors that, in principle, could affect the fit between language representations in humans and language models. For example, languages with templatic (non-concatenative) morphology, such as Hebrew, may make it challenging for standard sub-word tokenization approaches to capture their full morphological complexity (de Varda & Marelli, 2023; Klein & Tsarfaty, 2020). In turn, this mismatch between sub-word segmentation in models and speakers’ representations could affect the predictivity of the brain encoding models. Relatedly, languages where the written form underspecifies the content expressed in spoken form could pose additional challenges to text-based encoding models. For example, Hausa distinguishes grammatical classes through tone but lacks tone marking in its orthography (Adebara & Abdul-Mageed, 2022); thus, the language models’ input will inevitably lack some informative features which may contribute to brain alignment.

The English-centric approach to language hinders both cognitive science (Blasi et al., 2022) and NLP research (Joshi et al., 2020). Research at the intersection of these fields has struggled to widen its focus beyond English and provide a globally representative picture of the computational principles underlying language processing in humans. As a result, the generalizability of some of the claims in the computational cognitive science of language has been questioned, suggesting that the objective functions that neural network language models are trained to optimize might not be a suitable basis for language-general *in silico* models of human language processing (Blasi et al., 2022). Although these concerns are certainly warranted, our findings bring optimism to the field, demonstrating that many claims previously reported for English do, in fact, generalize across languages. This result suggests that the principles underpinning language processing in neural network language models are broadly applicable to the human language system, supporting the idea that these computational models capture fundamental aspects of human language processing.

## Methods

### Study I

#### fMRI experiments

Our first study utilized pre-existing functional magnetic resonance imaging (fMRI) data from Malik-Moraleda, Ayyash, et al. (2022) focusing specifically on brain responses elicited by the “passage listening” paradigm. In this setup, participants were exposed to passages approximately 4.5 minutes in duration, presented in their native language.

#### Participants

The original dataset included fMRI recordings from 86 right-handed participants, with a balanced gender distribution (50% females), ranging in age from 19 to 43 years (mean = 27.52, SD = 5.49). The dataset encompassed 45 languages across 12 language families. For 41 of these languages, there were two participants each, whereas the remaining four languages were represented by just one participant each. The number of participants considered in our study was subsequently reduced based on the reliability assessment of the fMRI responses (see Methods, “Study II – Time series extraction”). The languages with only one participant were not considered in our study due to the inability to establish response reliability.

#### Critical fMRI task

In Malik-Moraleda, Ayyash, et al.’s (2022) study, participants engaged in a variety of tasks while undergoing fMRI scanning. These tasks included a standard language localizer in English (all participants were proficient English speakers), a critical localizer in the participants’ native languages, two non-linguistic tasks, a resting-state scan, and a naturalistic passage listening task. Our analyses focused on the passage listening task, where participants listened to ∼4.5 min long passages from the book *Alice in Wonderland* (Carroll, 1865), recorded by a female native speaker of each represented language. Because the passage recordings varied slightly in length across languages, they were all trimmed to 260 s, and the final two seconds of the recording included a gradual volume fade-out. In addition, each run included 12 seconds of silence at the beginning and end, for a total duration of 284 s (4 min 44 s).

#### fMRI data acquisition

Structural and functional data were collected on the whole-body 3 Tesla Siemens Trio scanner with a 32-channel head coil at the Athinoula A. Martinos Imaging Center at the McGovern Institute for Brain Research at MIT. T1-weighted structural images were collected in 179 sagittal slices with 1 mm isotropic voxels (TR = 2,530 ms, TE = 3.48 ms). Functional, blood oxygenation level dependent (BOLD) data were acquired using an EPI sequence (with a 90° flip angle and using GRAPPA with an acceleration factor of 2), with the following acquisition parameters: thirty-one 4mm thick near-axial slices, acquired in an interleaved order with a 10% distance factor; 2.1 mm × 2.1 mm in-plane resolution; field of view of 200mm in the phase encoding anterior to posterior (A >> P) direction; matrix size of 96 × 96; TR of 2000 ms; and TE of 30 ms. Prospective acquisition correction was used to adjust the positions of the gradients based on the participant’s motion one TR back. The first 10 s of each run were excluded to allow for steady-state magnetization.

#### fMRI data preprocessing

Data preprocessing was performed with SPM12 (using default parameters, unless specified otherwise) and custom scripts in MATLAB. Preprocessing of anatomical data included normalization into a common space (Montreal Neurological Institute [MNI] template) and segmentation into probabilistic maps of the gray matter (GM), white matter (WM), and cerebro-spinal fluid (CSF). A GM mask was generated from the GM probability map and resampled to 2 mm isotropic voxels to mask the functional data. The WM and CSF maps were used as described below. Preprocessing of functional data included motion correction (realignment to the mean image using second-degree b-spline interpolation) and normalization into the MNI space (estimated for the mean image using trilinear interpolation). For this step, we used SPM12’s unified segmentation and normalization procedure with a reference functional image computed as the mean functional data after realignment across all timepoints omitting outlier scans. The output data were then resampled to a common bounding box between MNI-space coordinates (−90, −126, and −72) and (90, 90, and 108), using 2 mm isotropic voxels and 4th degree B-spline interpolation. Finally, the functional data were smoothed spatially using spatial convolution with a 4 mm full-width half-maximum (FWHM) Gaussian kernel.

#### fMRI data modeling and definition of functional regions of interest (fROIs)

The language and multiple demand network localizer data (here and in Study II) as well as the data from the additional training datasets in Study II and III were all modeled using the same setup. Effects were estimated using a General Linear Model (GLM) in which each experimental condition was modeled with a boxcar function convolved with the canonical hemodynamic response function (HRF) (fixation was modeled implicitly, such that all timepoints that did not correspond to one of the conditions were assumed to correspond to a fixation period). Temporal autocorrelations in the BOLD signal timeseries were accounted for by a combination of high-pass filtering with a 128 seconds cutoff, and whitening using an AR(0.2) model (first-order autoregressive model linearized around the coefficient a=0.2) to approximate the observed covariance of the functional data in the context of Restricted Maximum Likelihood estimation (ReML). In addition to experimental condition effects, the GLM design included first-order temporal derivatives for each condition (included to model variability in the HRF delays), as well as nuisance regressors to control for the effect of slow linear drifts, subject-motion parameters, and potential outlier scans on the BOLD signal.

Language fROIs were defined in individual participants (Saxe et al., 2006; Fedorenko et al., 2010) by combining two sources of information: (1) the participant’s activation map for the localizer contrast (Fedorenko et al., 2010) and (2) group-level constraints (‘parcels’) that delineated the expected gross locations of activations and were sufficiently large to encompass the variability in the locations of individual activations. The primary language localizer, described in detail in Fedorenko et al. (2010), consisted of reading English sentences and lists of unconnected nonwords in a standard blocked design, with stimuli presented one (non)word at a time. The parcels were derived from a probabilistic activation overlap map using watershed parcellation, as described by Fedorenko et al. (2010), for the *sentences > non-words* contrast in 220 independent participants and covered extensive portions of the lateral frontal, temporal, and parietal cortices. Five language fROIs were defined (as the 10% of voxels with the highest t-values for the *sentences > nonwords* contrast) in the dominant hemisphere (left for all participants but one Marathi speaker): three on the lateral surface of the frontal cortex and two on the lateral surface of the temporal and parietal cortex.

We used the English localizer in our main analyses to ensure comparability with previous studies and because it has been established that the localizer for a language works well as long as participants are fluent in that language (Chen et al., 2021; Malik-Moraleda, Ayyash et al., 2022). For completeness, we also used a native-language auditory localizer (Malik-Moraleda, Ayyash et al., 2022), contrasting intact passages with acoustically degraded passages; results were consistent across the two localizers (SI S7).

To test for hemispheric specificity, we additionally defined five homotopic language fROIs in the right hemisphere. As a further control, we examined fROIs in the multiple demand (MD) network, a bilateral set of areas implicated in executive functions and fluid intelligence (Duncan, 2010; Fedorenko et al., 2013; Assem et al., 2020a). The MD fROIs were defined using a spatial working memory localizer task (e.g., Assem et al. 2020b), following the same participant-specific, parcel-constrained procedure as for the language network. Parcels for the MD network were derived from a large independent sample (Assem et al., 2020b). All parcels are available for download (https://www.evlab.mit.edu/resources-all/download-parcels).

#### Time series extraction

The passages were 260 s (4 min 20 s) long, surrounded by 12-s periods of silence. fMRI responses were recorded at the temporal resolution of 2 s, resulting in 142 data points (volumes or TRs) (130 TRs after the exclusion of the silence periods). For each participant, BOLD time series were extracted from each voxel within the left-hemisphere language network, as well as their right-hemisphere homologues, and the multiple demand (MD) network. Within each system, voxel-wise responses were averaged to obtain one time series per fROI. In addition, we considered the average response across the five language fROIs in the left hemisphere, and analogously across the right-hemisphere and MD fROIs. To account for the lag in the hemodynamic response, we shifted the time series by 6 seconds relative to the linguistic input (i.e., the word embeddings aligned to the stimulus). This shift ensured the appropriate temporal alignment between the stimulus representations and the brain responses for the encoding analyses (this choice was corroborated by encoding analyses in a separate dataset; SI S9). As noted above, we focused on the subset of languages where the language network time series of the two participants showed a positive and statistically significant Pearson correlation (*p* < .05; see SI S10 for the results of a sanity-check analysis showing that the correlations are higher within a language than between languages). This selection criterion left 12 languages: Afrikaans, Dutch, Farsi, French, Lithuanian, Marathi, Norwegian, Romanian, Spanish, Tamil, Turkish, Vietnamese (see **Figure 4A**), excluding 29 languages. We note that the reliability of the time series varied substantially across the 12 languages we retained (ranging from *r* = 0.196 for Marathi to *r* = 0.462 for French; Figure 4A), and we examine the consequences of this variability for cross-lingual transfer in SI S14.

**Figure 4.**
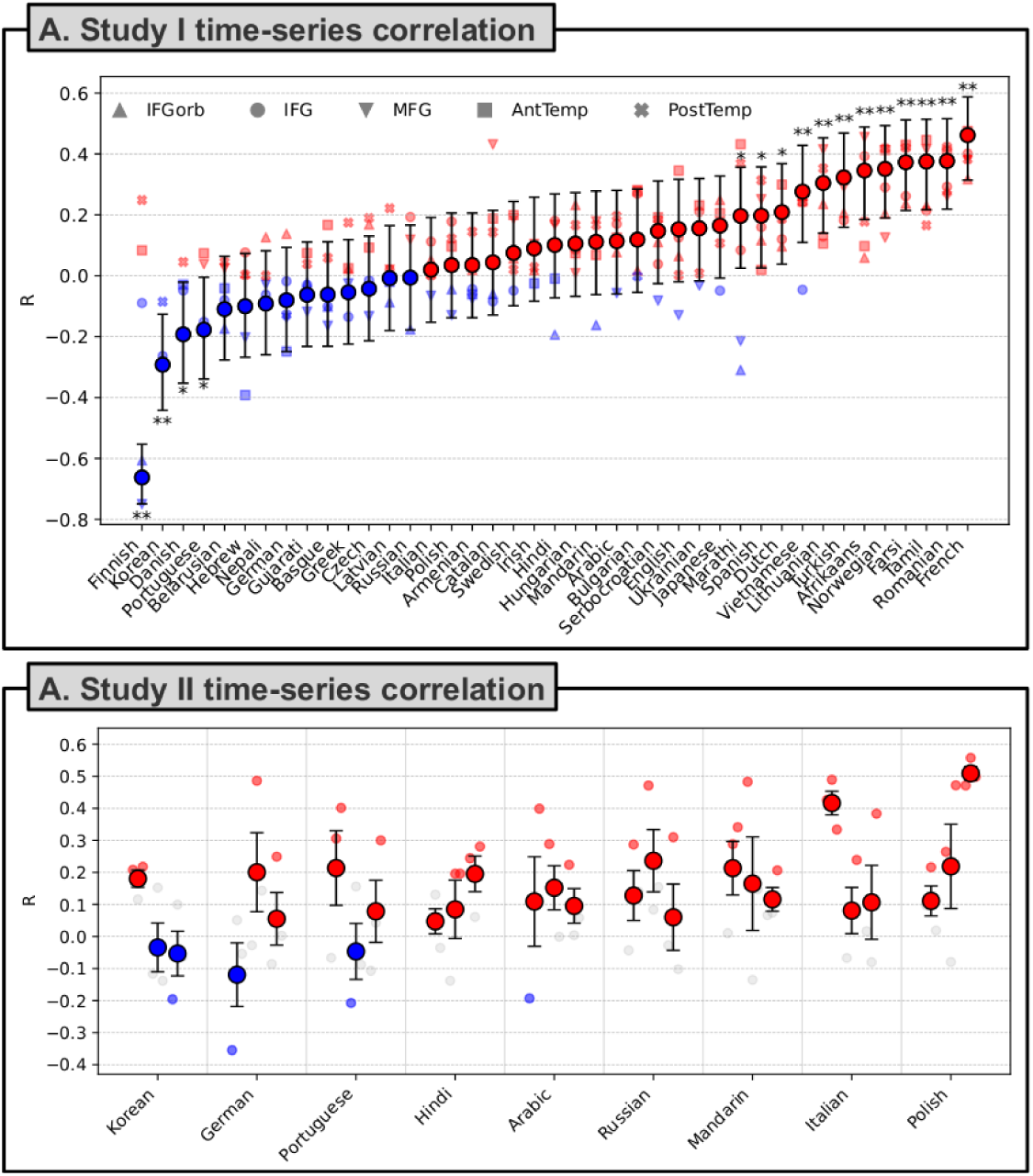
fMRI time series data reliability and temporal alignment of the embeddings with the time series. **A. Study I time series correlation:** Correlation between the time series of the two participants for each language (Study I data, from Malik-Moraleda, Ayyash, et al., 2022). The large dots indicate the correlations in the language network, averaging the time series across fROIs, whereas the smaller markers indicate the correlations in the five language fROIs. The color of the dots indicate whether the correlation is positive (red) or negative (blue). The asterisks indicate statistical significance (*p* < .05*; *p* < 0.01**). **B. Study II time series correlation:** Correlation between the time series of the three participants for each language × passage combination (Study II data). For each language, we have 3 large dots: each dot represents the correlation in the time series in response to one of the 3 passages, averaging across participant pairs. Three smaller markers are associated with each large dot, showing the correlations for all possible participant pairs (participants 1 and 2, 1 and 3, and 2 and 3).

#### Timestamped speech recognition

Participants were exposed to continuous auditory stimuli. To convert these audio stimuli into textual data suitable for language model analysis, we employed an automated transcription process using the Whisper-timestamped tool (Louradour, 2023). This tool is an extension of the Whisper model (Radford et al., 2023) and it outputs precise word-by-word timestamps. We utilized the largest available model variant (1,550M parameters); this model supported transcription in all the selected languages.

#### Language models

Following the acquisition of the textual input associated with each passage, we generated contextualized text embeddings using a variety of MNNLMs. The specific models employed are detailed in Table 1 and feature a range of architectures, including encoder-only, decoder-only, and encoder-decoder models, with parameter counts ranging from 118 million to 4.5 billion. Some of the models were explicitly optimized to align representations across languages; some examples include models trained on translation (NLLB), word alignment in translation pairs (XLM-Align), and contrastive cross-lingual training (InfoXLM). The majority of the models, however, were not trained with an explicit cross-lingual objective; instead, their pre-training tasks involved the reconstruction of the input text that had been altered or corrupted in some way. The models trained on masked language modeling (mBERT, XLM-R) were optimized to predict randomly masked tokens within the input by relying on bidirectional context, and the models based on span corruption (mT5) were similarly trained to reconstruct consecutive spans of text; mDeBERTa was trained to detect tokens that had been replaced in the input. Causal language models (mGPT, XGLM) were trained to predict each token sequentially based on the preceding context only. We also considered two models (DistilmBERT, mMiniLM) trained by distilling the language knowledge from larger, more complex masked language models (mBERT, XLM-R) into smaller, more efficient ones. All models were utilized as-is, without any fine-tuning or adaptation, and were accessed via the transformers Python package (version 4.45.2; Wolf et al., 2020).

We extracted the hidden states from each layer of each MNNLM in response to the transcribed text and derived word-level embeddings by averaging the embeddings of sub-word tokens (see “Encoding models” section below for procedure in case words were split between TRs). In the case of encoder-decoder models, we did not consider the representations generated by the decoder. We added model-specific special tokens (e.g., beginning-of-sequence and end-of-sequence tokens) to the input text where appropriate, and then discarded their corresponding embedding representations.

All the models in our sample except the ones based on a decoder-only architecture were bidirectional, meaning that token representations could incorporate information about future tokens. To prevent the models from accessing future information, we iteratively slid a window of 100 words across the texts, ensuring that each token’s embedding was influenced only by the preceding context within the window. This method allowed us to approximate a causal setup in bidirectional models, making the processing scheme more comparable with human language processing, where comprehension unfolds incrementally based on prior context without knowledge of future input. In SI S11, we considered an alternative approach of obtaining contextualized embeddings where we reset the context at each cross-validation fold boundary, ensuring that the embeddings could not carry over contextual information from one fold to the next. This analysis showed that preventing context bleed-over had a limited effect on performance.

To assess how similar the training and test materials were, we computed an embedding-based similarity measure: for each training dataset, we calculated the cosine similarity between every word embedding in that dataset and every word embedding in the test dataset, and then averaged across words, sentences/passages, languages, layers, and models (**Figure 1C**).

#### Brain encoding models

We aligned the embeddings generated from the MNNLMs with the brain responses of participants exposed to the same linguistic stimuli. For each of the 130 TRs, TR-level embeddings were computed by averaging the embeddings of the words pronounced within that interval. Words that were pronounced at the TR boundaries (e.g., a word whose pronunciation started in a given TR and ended in the subsequent one) were included in the embedding calculation of the second TR. We normalized with *z*-scoring both the embedding representations (per unit) and the fMRI time series (per time series). The normalization parameters were estimated in the training set and directly applied to the test set. The mapping of the embeddings to brain responses was accomplished using ridge (i.e., *L2*-penalized) regression. The α parameter, which controls the strength of the regularization, was set in an inner loop with nested five-fold cross-validation by considering 10 candidate values ranging from 10^−5^ to 10^4^, logarithmically spaced. This was achieved with the RidgeCV function of the package himalaya (version 0.4.6; La Tour et al., 2022).

### WITHIN encoding models

As a first step of our study, the encoding models were trained and tested on non-overlapping data subsets within each language. In these encoding models, the predictor variable was each TR-level embedding representation, whereas the dependent variable was each participant’s response in each of the five language fROIs of interest, averaged across voxels, as well as the average response across all five fROIs. The same procedure was adopted for the right-hemisphere fROIs and the 20 fROIs of the Multiple Demand network. Critically, the encoding models were evaluated across participants, such that a model was fit with 10-fold cross-validation on 90% of the data from one participant, and evaluated on the remaining 10% of the data from the other participant. Data from each of the two participants was used separately for training and testing (i.e., training on participant A’s data and testing on participant B’s data, then vice versa), and the results were then averaged. The encoding performance was assessed via 10-fold cross-validation with sequential splits^1^. We adopted a sequential train-test split since with shuffled train-test splits encoding models can achieve high predictivity by simply relying on the temporal autocorrelation of the signal (Feghhi et al., 2024; Kauf, Tuckute et al., 2024). The predicted responses for each left-out fold were then concatenated to reconstruct a full time series, which was then compared to the recorded brain responses via Pearson correlation. We developed separate encoding models for every combination of fROI, language, model, and layer. Note that although most models were fit for all the 12 languages considered in the study (see SI S4 for data from the 9 languages from Study II), the encoding models based on the XGLM embeddings were fit in 5 languages (Spanish, Vietnamese, Tamil, Turkish, French) and the encoding models based on mGPT were fit in 10 languages (Afrikaans, Farsi, French, Lithuanian, Marathi, Romanian, Spanish, Tamil, Turkish, Vietnamese), as the remaining languages were absent from the models’ pre-training data. A graphical summary of the encoding procedure is provided in **Figure 1B**, top.

### ACROSS encoding models

In a subsequent step, we tested whether the encoding models could be transferred to new languages. To this end, for each individual language *li* ∈ L, we fit an encoding model in all the remaining languages *lj* ∈ L \ {*li*}—that is, concatenating 90% of the data from participant pairs listening to all but the target language. Then, the obtained regression weights were employed to predict the remaining 10% of the fMRI responses in *li*, which was not included in the training data. This procedure was repeated with cross-validation until the full brain responses in *li* were predicted. Then, the encoding performance was evaluated as the Pearson correlation between the predicted and the actual brain response in *li*, calculated separately for each participant belonging to *li*, and then averaged across the two participants. In a supplementary analysis, we also trained the encoding models in a single language *li* ∈ L and evaluated them in all the remaining languages *lj* ∈ L \ {*li*}. While the raw result scores were lower due to the smaller amount of training data, encoding models trained in a single language generalized robustly across languages (SI S14). A schematic visualization of the cross-lingual encoding models is displayed in **Figure 1B**, bottom.

#### Significance testing

To evaluate the statistical significance of the encoding performance scores, we compared the results of the encoding models with the scores obtained by equivalent encoding models where the fMRI responses underwent a random circular shift (i.e., destroying the temporal correspondence between stimulus and brain response while keeping the temporal autocorrelation of the signal intact). Specifically, we circularly shifted the fMRI time series by equally spaced offsets *o* ∈ {26, 52, 78, 104} TRs and re-ran the same 10-fold encoding analysis for each of those values of *o* separately.

Statistical significance was evaluated via a two-step analytical approach measuring whether the correlations obtained by the encoding model were systematically higher than the ones obtained after shifting the response variable. First, we calculated the difference between corresponding pairs of Pearson correlation coefficients from the original and the shifted encoding models, where each pair of coefficients corresponds to the scores obtained by a given encoding model in a specific language. Given the non-normal distribution of Pearson correlation scores, we applied Fisher’s *r*-to-*z* transformation to convert these coefficients into *z*-scores (Fisher, 1921; see Eq. 1). The difference between the obtained *z-*scores was divided by the standard error of the difference, i.e., 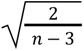, where *n* = 130 is the sample size used to calculate the correlation coefficient. This procedure allowed us to generate a *z*-statistic for each correlation pair, reflecting the magnitude and direction of the difference between the correlation coefficients obtained by the original and the randomized encoding models.

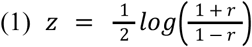

To assess the overall significance across all pairs of correlations obtained in the various languages by a given MNNLM, we combined the individual *z*-statistics derived in the previous step using Stouffer’s method (Stouffer et al., 1949), which is more conservative than Fisher’s combined probability method in that it does not assume independence between observations. This method involves summing the *z*-statistics and dividing by the square root of *k*, the number of statistics combined (Eq. 2).

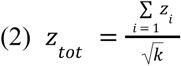

The combined *z_tot_* statistic was then converted into a two-tailed *p*-value, reflecting the probability of observing a combined difference in correlations as extreme as, or more extreme than, the one observed given a null hypothesis. This significance calculation procedure was carried out independently for each encoding model, both in the context of WITHIN and ACROSS encoding models.

#### Next-word prediction performance

To assess whether next-word prediction performance explained how well the model representations fit to the brain data, we evaluated all models on a word prediction task using a custom multilingual dataset. To create this dataset, we first scraped 50,000 words from random Wikipedia pages in each of the 12 languages considered in the study (6M words in total). We removed titles and headings and considered only text paragraphs. Since meaningful comparisons in next-word prediction ability can be drawn only in the context of similarly sized vocabularies, we defined a vocabulary in each language as the 5,000 most common words in that language. Then, we divided the continuous text streams into chunks of 10 words each by sliding a 10-words window on the text data.

Next, we extracted word embeddings from all 20 language models considered in this study; for each 10-word chunk, we considered the contextualized embeddings of the last word, as derived from the last layer of the model. Lastly, we trained a ridge multi-class classifier to predict the following word (encoded as a one-hot 5,000-dimensional vector) from the contextualized embedding of the previous one. As for the neural encoding models, the α parameter was set with nested cross-validation. We trained the classifier independently in each language with stratified 10-fold cross-validation, to ensure that all classes were present in both the training and the test set. We evaluated the models’ next-word prediction ability as the perplexity on held-out data, averaged over folds. Perplexity is a standard metric in language modeling that reflects how well a model predicts the correct next word: lower values indicate that the model assigns higher probability to the observed word, corresponding to better predictive performance. Random seeds for all random processes in the pipeline were set to ensure reproducibility. In preliminary analyses, we found that the perplexity values obtained by the models in each language were highly correlated with the perplexity values obtained by a randomized baseline, possibly due to the fact that some languages are more predictable than others due to different vocabulary sizes and word repetition rates. To isolate the next-word prediction abilities of the MNNLMs, we used the perplexity estimates generated by the randomized models to linearly predict the models’ actual perplexity estimates, and used as a next-word prediction metric the residual of the prediction, effectively decorrelating the target estimates with the ones produced by the randomized models. Similar results were obtained using the raw perplexity estimates. The relationship between next-word prediction scores and encoding performance was assessed as the Pearson correlation between each model’s residual perplexity values and its best-layer encoding score obtained in the WITHIN and ACROSS conditions.

### Study II

#### fMRI experiments

Similar to Study I, our second study was based on fMRI data from participants listening to passages in their native language. In this setup, participants were exposed to three passages approximately 4.5 minutes in duration. All the methodological details (fMRI data acquisition, preprocessing, definition of fROIs) were similar to Study I except that the MRI scanner had undergone a Prisma upgrade (see Methods, “Study II – fMRI data acquisition”).

The study was performed with ethical approval from the Committee on the Use of Humans as Experimental Subjects at the Massachusetts Institute of Technology (MIT) (protocol number 2010000243). All participants gave informed written consent before starting the experiments.

#### Participants

We recruited 27 participants (three in each of the 9 target languages, i.e., Italian, Portuguese, German, Russian, Polish, Hindi, Arabic, Mandarin, Korean). All but two participants were right-handed; the remaining two were left-handed and ambidextrous; participants ranged in age from 20 to 42 years (mean = 27.37, SD = 5.18), and 18 of them were females (66.66%). None of the participants had taken part in Malik-Moraleda, Ayyash, et al.’s (2022) study, so there was no overlap in participants between Study I and Study II.

#### Critical fMRI task: Passage listening

The critical fMRI task was equivalent to the one in Study I, except that participants listened to three passages (as opposed to one) in their native language. The passages all came from the book *Alice in Wonderland* (Carroll, 1865) and were recorded by a female native speaker; the same criteria were applied with respect to the passages’ durations.

#### fMRI data acquisition

Structural and functional data were collected on a whole-body 3 Tesla Siemens Prisma scanner with a 32-channel head coil at the Athinoula A. Martinos Imaging Center at the McGovern Institute for Brain Research at MIT. T1-weighted, Magnetization Prepared Rapid Gradient Echo (MP-RAGE) structural images were collected in 208 sagittal slices with 1 mm isotropic voxels (TR = 1,800 ms, TE = 2.37 ms, TI = 900 ms, flip = 8 degrees). Functional, blood oxygenation level-dependent (BOLD) data were acquired using an SMS EPI sequence with a 90° flip angle and using a slice acceleration factor of 3, with the following acquisition parameters: seventy-two 2 mm thick near-axial slices acquired in the interleaved order (with 10% distance factor), 2 mm × 2 mm in-plane resolution, FoV in the phase encoding (A >> P) direction 208 mm and matrix size 104 × 104, TR = 2,000 ms, TE = 30 ms, and partial Fourier of 7/8. The first 10 s of each run were excluded to allow for steady state magnetization.

#### Time series extraction

As in Study I, language fROIs were defined using the standard English functional localizer (Fedorenko et al., 2010). The passages were 260 s long, surrounded by 12-s periods of silence. With a temporal resolution of 2 s, each passage contributed 142 data points (130 after the exclusion of the silence periods). For each participant, BOLD time series were extracted from each voxel within the language network; then, they were averaged across the voxels in each of the five language fROIs, and then across fROIs; lastly, they were averaged across participants (two or three, depending on the correlations in the time series), unlike Study I, where participant data were analyzed separately. For each language, correlations between the participants’ time series were assessed across the three passages. **Figure 4C** displays the correlations between participant pairs within each passage for each language. Although some individual language × passage combinations displayed near-zero or negative correlations, the correlations were generally positive and significant across languages. In line with our approach in Study I, we retained only participant pairs with time series that were significantly correlated at the level of each individual passage. Importantly, this procedure allowed for variation in participant pairs across passages within the same language. For instances where two out of three participant pairs showed significant correlations in their time series, all three participants were included, since all three time series correlated with at least one of the others. Using these criteria, we obtained reliable time series for all three passages in each language, with the exception of five specific language-passage combinations: Korean (passages 2 and 3), German (passage 1), Portuguese (passage 2), and Hindi (passage 1). SI S10 reports the results of a sanity-check analysis showing that the correlations in the time series are higher within a language than between languages.

#### Training data

In our second experiment, we fit several encoding models from different data sources and transferred them zero-shot to the newly collected data. The original data sources encompass different types of stimuli and modalities of presentation. In particular, in addition to the data employed in our first study, we considered two datasets where participants read sentences (*Pereira2018*, Pereira et al., 2018; *Tuckute2024*, Tuckute et al., 2024a), and one additional dataset where participants listened to stories in English (*NatStories*, expanded from Blank et al., 2014). Across all datasets in Study II, we extracted responses from the same five language fROIs (see “fMRI data modeling and definition of functional regions of interest (fROIs)”) and averaged them across fROIs, and then across participants, thus following the same procedure as in Study I.

#### Pereira2018

The *Pereira2018* dataset (Pereira et al., 2018; Exp. 2 and 3) includes fMRI recordings from 10 unique participants (9 in Exp. 2 and 6 in Exp. 3) reading 627 sentences (384 in Exp. 2, 243 in Exp. 3), grouped into passages of three to four sentences. Each participant saw each sentence between 1 and 4 times, with the majority seeing each sentence 3 times. The two sets of materials were constructed independently, and each spanned a broad range of content areas. Sentences were 7–18 words long in Experiment 2, and 5–20 words long in Experiment 3. The sentences were presented visually on a screen one at a time for 4 seconds (followed by 4 seconds of fixation, with additional 4 seconds of fixation at the end of each passage).

#### Tuckute2024

The *Tuckute2024* dataset (Tuckute et al., 2024a) includes fMRI recordings from 10 unique participants reading 1,000 sentences. The materials were extracted from written and transcribed spoken corpora and were designed to maximize semantic content and stylistic diversity. All sentences were 6 words long. The sentences were presented visually on a screen one at a time for 2 seconds (followed by 4 seconds of fixation). Each participant read each sentence once, and data were acquired across two or three separate scanning sessions.

#### fMRI preprocessing (shared between Pereira2018 and Tuckute2024)

The same preprocessing pipeline was used as for the localizer data in Study I and Study II.

#### First-level Analysis (Pereira2018)

The same modeling setup was used as for the localizer data in Study I and Study II. Each sentence trial was modeled as a separate condition.

#### First-level Analysis (Tuckute2024)

Effects were estimated using a GLM in which each experimental condition (i.e., each sentence trial) was modeled using GLMsingle (Prince et al., 2022). Using this framework, a general linear model (GLM) was used to estimate the beta weights that represent the BOLD response amplitude evoked by each individual sentence trial (fixation was modeled implicitly, such that all time points that did not correspond to one of the conditions (sentences) were assumed to correspond to a fixation period). For each voxel, the HRF that provided the best fit to the data was identified (on the basis of the amount of variance explained). The data were modeled using a fixed number of noise regressors (five) and a fixed ridge regression fraction (0.05) (these parameters were determined empirically using a joint data modeling and data evaluation framework; see Tuckute et al., 2024a). By default, GLMsingle returns beta weights in units of percent signal change by dividing by the mean signal intensity observed at each voxel and multiplying by 100. To mitigate the effect of collecting data across multiple scanning sessions, the beta values were *z*-scored session-wise per voxel (Methods, “fMRI data modeling and definition of functional regions of interest (fROIs)”).

#### NatStories

The *NatStories* dataset is a collection of fMRI recordings from participants listening to naturalistic stories (∼5 minutes each) while in the fMRI scanner. The dataset was collected in the Fedorenko lab between 2014 and 2020. The first findings were reported in Blank et al. (2014), and the data from the expanded dataset were used in several other studies (Paunov et al., 2022; Shain, Blank, et al., 2020; Shain et al., 2022; Sueoka, Paunov et al., 2024; Wehbe et al., 2021), but the dataset has never been used in its entirety, as we do here. The dataset features 9 stories: 8 were edited to include low-frequency syntactic constructions but maintaining fluency for native speakers (see Futrell et al., 2018 for details of these materials), and 1 was an expository text (“Tree”) adapted from Wikipedia (see Paunov et al., 2022 for details). The number of participants per story varied from 7 to 70 (see SI S12). Because this is the first time the dataset is used in its current form, we conducted a reliability analysis which showed that the time series were highly reliable (see SI S12).

#### fMRI preprocessing (NatStories)

The same preprocessing pipeline was used as for the localizer data in Study I and Study II. Additional preprocessing of data was performed using the CONN toolbox (Whitfield-Gabrieli & Nieto-Castañón, 2012; https://www.nitrc.org/projects/conn) with default parameters, unless specified otherwise. BOLD signal time courses were extracted from WM and CSF. Five temporal principal components were obtained from each, as well as voxel-wise averages. These were regressed out of each voxel’s time course, along with additional noise regressors, specifically, six motion parameters estimated during offline motion correction (three translations and three rotations) and their first temporal derivatives, and artifact time points (based on global signal outliers and motion).

#### Brain encoding models

As in Study I, our main analyses were based on the training and evaluation of encoding models predicting brain activity from MNNLM-based embeddings. The general procedure that allowed us to derive TR-level embeddings was left unaltered with respect to Study I, and involved obtaining speech transcriptions with Whisper-timestamped, computing the embeddings, and temporally aligning the embeddings with the time series. Differently from Study I, we here considered a single layer for each MNNLM: the layer that we identified in Study I as the best-performing layer in the ACROSS condition. All the languages we considered in Study II were present in the models’ pre-training data.

We fit several encoding models from different data sources (one for each model × data source combination) and transferred them zero-shot to the new data we collected. In the case of sentence-level datasets (*Pereira2018*, *Tuckute2024*), the embedding representations for the complete sentences were obtained by averaging over all the words in the sentence; thus, each sentence contributed to a single training example for the encoding models. In the case of the naturalistic passages (Study I data, *NatStories*), we aligned the embeddings with the fMRI time series employing the same procedure as in Study I, averaging the embedding representations over the relevant TRs. All the technical aspects of the encoding models (e.g., penalty estimation, normalization), were left unaltered from Study I. Statistical significance was assessed against randomized baselines designed to preserve dataset-specific structure: for continuous passage data (Study I data, *NatStories*), fMRI responses were circularly shifted by 26, 52, 78, or 104 TRs to preserve the temporal autocorrelation of the signal, while for sentence-level data (*Pereira2018*, *Tuckute2024*), the correspondence between sentences and responses was randomly shuffled, repeating this procedure four times with four different random seeds. Encoding scores were aggregated first across passages, then across languages.

#### Similarity in the language materials

Unlike Study I, where the language materials consisted of passages from the same book and thus were highly homogeneous, Study II involved the transfer of encoding models across very different kinds of linguistic stimuli. To assess the similarity between the linguistic materials used for training the encoding models (Study I materials, *NatStories*, *Pereira2018*, *Tuckute2024*) and those used for testing them (Study II materials), we considered both the similarity in the word embeddings produced by the MNNLMs in our sample and the word overlap across the stimuli. To calculate embedding-based similarity, for each training dataset, we computed the cosine similarity between each word embedding from the dataset’s materials and each word embedding in the test dataset. Similarity values were then averaged across words, sentences/passages, languages, layers, and models (**Figure 1C**). To measure word overlap, we calculated the number of unique words shared between each training dataset and the test dataset (Wtrain ∩ Wtest) divided by the number of unique words included in both datasets (Wtrain ∪ Wtest). Their ratio, which corresponds to the Jaccard index, was employed as a metric of lexical overlap.

### Study III

We constructed perturbed versions of the English stimuli following Kauf, Tuckute, et al. (2024). We considered three manipulation types: “information loss” manipulations that removed subsets of lexical categories, word-order manipulations that disrupted the sequential structure of the sentences, and paraphrase manipulations that changed the sentences’ surface form while preserving meaning.

Information loss perturbations were based on part-of-speech (POS) tagging using the NLTK perceptron tagger (Bird & Loper, 2004). Six variants were generated: sentences retaining only (i) content words (nouns, verbs, adjectives, adverbs), (ii) nouns, verbs, and adjectives, (iii) nouns and verbs, (iv) nouns only, (v) verbs only, or (vi) function words only. Perturbed sentences were created by removing all tokens not matching the relevant POS categories.

Word order was manipulated in two ways. First, local word swaps exchanged adjacent tokens, with between 1 and 5 swaps applied per sentence; swap indices were selected with a seeded random generator to ensure reproducibility. Second, a full reversal condition inverted word order in each sentence, so that the sentence was presented from the last word to the first one.

Paraphrases were generated with the Llama-4 Maverick 17B-128E model (accessed via the Together API), using greedy decoding (temperature = 0). Outputs were manually inspected and corrected in a small number of cases (e.g., formatting errors or incomplete outputs).

For each perturbation type, we extracted embeddings from the set of MNNLMs considered throughout Study I and Study II. For each model we identified the best layer based on the results from Study I. Encoding models were trained to predict fMRI responses in the perturbed versions of the English stimuli of the *Pereira2018* and *Tuckute2024* datasets. All the details regarding the fit of the encoding models was left unaltered with respect to Study II. We tested whether each perturbation reduced encoding relative to the intact condition by computing Fisher *r*-to-*z* differences for each model × dataset × language × passage combination, then aggregating *z*s with Stouffer’s method (two-tailed *p*) and applying Bonferroni correction across perturbation types.

### Study IV

Study IV complements the stimulus perturbation approach used in Study III with an analysis that operates directly on the embedding representations. We applied Iterative Null-space Projection (INLP; Ravfogel et al., 2020) to selectively remove meaning-related or form-related information from the MNNLM embeddings, and then evaluated the effect of these forms of ablation on cross-lingual transfer (see SI S17 for a comparison with results within a single language).

#### Features

We defined two feature sets to characterize meaning- and form-related properties of the sentences in the *Tuckute2024* dataset. The meaning feature set includes human ratings on seven dimensions, collected by Tuckute et al. (2024a): imageability, others’ thoughts, physical interaction, places, valence, arousal, and plausibility. The form feature set also includes seven features: PCFG negative log-probability (surprisal), grammaticality ratings, overall frequency ratings, and conversational frequency ratings from the same dataset, plus three automatically computed features, namely mean dependency length (computed with the spaCy English pipeline; Honnibal et al., 2020), mean word frequency (Zipf scale), and mean word length in characters. All features were computed at the sentence level for the 1,000 baseline sentences in the *Tuckute2024* dataset.

#### Iterative Null-space Projection (INLP)

Having obtained embeddings from the models with the same procedure described for Studies I–III, we applied a regression-based variant of INLP (originally developed for classification; Ravfogel et al., 2020) to remove linear information about each feature set from the embeddings.

The procedure is composed of the following steps:

1. Standardize the embedding matrix *X* and feature matrix *Y*;
2. Fit a ridge regression model predicting one target feature from the embeddings;
3. Collect the regression weights as a vector, which defines the direction in the embedding space that is most predictive of that feature;
4. Compute a projection that maps the embeddings onto the null space of all weight vectors collected so far (that is, onto the subspace of the embedding space that is orthogonal to all identified predictive directions) with SVD (using the Ben-Israel numerically stable formula as in Ravfogel et al., 2020);
5. Project the embeddings onto this null space, effectively removing the identified predictive information, and return to step (2) for the next feature.

The loop over the features was repeated until the maximum R^2^ across all features was below 0.01. At that point, the accumulated weight vectors were assembled into a matrix *W*, and the final null-space projection matrix *P* was computed via SVD. The final ablated embeddings were then obtained as *X_ablated_ = X · P*.

To assess the selectivity of the ablation, after removing the meaning features, we evaluated whether we could decode form features from the ablated embeddings, and vice versa. This analysis confirmed that INLP preferentially removed the targeted information, although the non-targeted features showed some degree of degradation as well, due to non-independence between the two feature sets.

#### Encoding model training and evaluation

For each MNNLM in our sample, we trained three encoding models on the *Tuckute2024* data: one using intact embeddings, one using meaning-ablated embeddings, and one using form-ablated embeddings. Encoding models were trained with the same procedure described for Studies II–III. At test time, word-level embeddings for the new languages were projected through the same null-space projection matrix *P* before temporal alignment with the fMRI time series, so that the same information was removed during both training and test. Statistical comparisons between the intact, meaning-ablated, and form-ablated conditions were performed with Fisher’s *r*-to-*z* transformation and Stouffer’s method for combining *z*-statistics across models and languages, following the same procedure described for Studies I–III.

## Data and code availability

Code and data to reproduce this study are available at https://osf.io/uc6y7/.

## Acknowledgements

We are grateful to Hee So Kim and Agata Wolna for assisting with participant recruitment; to Megha Vemuri, Alexander Fung, and Sara Swords for their assistance with data collection; and to Ben Lipkin for help with preprocessing and curating the Natural Stories dataset. AGdV was supported by the K. Lisa Yang ICoN Center Postdoctoral Fellowship. GT was supported by a Friends of McGovern Graduate Fellowship and the K. Lisa Yang ICoN Center Graduate Fellowship; EF was supported by NIH award U01-NS121471 and funds from the McGovern Institute for Brain Research, the Brain and Cognitive Sciences Department, the Simons Center for the Social Brain, and the MIT Quest for Intelligence.

## Contributions

**AGdV**: Conceptualization, Methodology, Investigation, Data curation, Visualization, Formal analysis, Validation, Software, Writing-Original draft; **SMM**: Investigation, Data curation, Writing-Review and editing; **GT**: Investigation, Data curation, Writing-Review and editing; **EF**: Conceptualization, Writing-Review and editing, Supervision, Project administration.

## Footnotes

1 In SI S11, we report the encoding performance obtained in a stricter condition where we reset the models’ context at each fold boundary to control for possible context contamination. Our results showed that context contamination did not substantially impact the results.

## References

Adebara, I., & Abdul-Mageed, M. (2022). Towards Afrocentric NLP for African languages: Where we are and where we can go. In Proceedings of the 60th Annual Meeting of the Association for Computational Linguistics (Volume 1: Long Papers), 3814–3841. Dublin, Ireland: Association for Computational Linguistics.

Antonello, R., & Huth, A. (2024). Predictive coding or just feature discovery? An alternative account of why language models fit brain data. Neurobiology of Language, 5(1), 64–79.

Antonello, R., Vaidya, A., & Huth, A. (2024). Scaling laws for language encoding models in fMRI. Advances in Neural Information Processing Systems, 36.

Assem, M., Glasser, M. F., Van Essen, D. C., & Duncan, J. (2020a). A domain-general cognitive core defined in multimodally parcellated human cortex. Cerebral Cortex, 30(8), 4361–4380.

Assem, M., Blank, I. A., Mineroff, Z. A., Ademoğlu, A., & Fedorenko, E. (2020b). Activity in the fronto-parietal multiple-demand network robustly associated with individual differences in working memory and fluid intelligence. Cortex, 131, 1–16.

Aw, K. L., & Toneva, M. (2023). Training Language Models to Summarize Narratives Improves Brain Alignment. In Eleventh International Conference on Learning Representations.

Belinkov, Y., Durrani, N., Dalvi, F., Sajjad, H., & Glass, J. (2020). On the linguistic representational power of neural machine translation models. Computational Linguistics, 46(1), 1–52.

Berlin, B., & Kay, P. (1969). Basic Color Terms: Their Universality and Evolution. Berkeley & Los Angeles University of California Press.

Bird, S., & Loper, E. (2004). NLTK. In Proceedings of the ACL 2004 on Interactive poster and demonstration sessions (pp. 31-es). Association for Computational Linguistics.

Blank, I., Kanwisher, N., & Fedorenko, E. (2014). A functional dissociation between language and multiple-demand systems revealed in patterns of BOLD signal fluctuations. Journal of neurophysiology, 112(5), 1105–1118.

Blasi, D. E., Henrich, J., Adamou, E., Kemmerer, D., & Majid, A. (2022). Over-reliance on English hinders cognitive science. Trends in cognitive sciences, 26(12), 1153–1170.

Braga, R. M., DiNicola, L. M., Becker, H. C., & Buckner, R. L. (2020). Situating the left-lateralized language network in the broader organization of multiple specialized large-scale distributed networks. Journal of neurophysiology, 124(5), 1415–1448.

Buchweitz, A., Shinkareva, S. V., Mason, R. A., Mitchell, T. M., & Just, M. A. (2012). Identifying bilingual semantic neural representations across languages. Brain and language, 120(3), 282–289.

Campbell, L. (2008). Ethnologue: Languages of the world.

Caucheteux, C., Gramfort, A., & King, J. R. (2021). Disentangling syntax and semantics in the brain with deep networks. In International conference on machine learning (pp. 1336–1348). PMLR.

Caucheteux, C., & King, J. R. (2022). Brains and algorithms partially converge in natural language processing. Communications biology, 5(1), 134.

Carroll, L. (1865). Alice’s adventures in wonderland (1st). Macmillan.

Chang, T. A., Arnett, C., Tu, Z., & Bergen, B. K. (2024). Goldfish: Monolingual Language Models for 350 Languages. arXiv preprint arXiv:2408.10441.

Chee, M. W., Caplan, D., Soon, C. S., Sriram, N., Tan, E. W., Thiel, T., & Weekes, B. (1999). Processing of visually presented sentences in Mandarin and English studied with fMRI. Neuron, 23(1), 127–137.

Chen, C., Gong, X., Tseng, C., Klein, D., Gallant, J., & Deniz, F. (2024). Bilingual language processing relies on shared semantic representations that are modulated by each language. bioRxiv, 2024-06.

Chen, X., Affourtit, J., Ryskin, R., Regev, T. I., Norman-Haignere, S., Jouravlev, O., … & Fedorenko, E. (2021). The human language system does not support music processing (p. 2021.06. 01.446439). bioRxiv.

Chi, E. A., Hewitt, J., & Manning, C. D. (2020). Finding Universal Grammatical Relations in Multilingual BERT. In Proceedings of the 58th Annual Meeting of the Association for Computational Linguistics. Association for Computational Linguistics.

Chi, Z., Dong, L., Zheng, B., Huang, S., Mao, X. L., Huang, H. Y., & Wei, F. (2021a). Improving Pretrained Cross-Lingual Language Models via Self-Labeled Word Alignment. In Proceedings of the 59th Annual Meeting of the Association for Computational Linguistics and the 11th International Joint Conference on Natural Language Processing (Volume 1: Long Papers) (pp. 3418–3430).

Chi, Z., Dong, L., Wei, F., Yang, N., Singhal, S., Wang, W., … & Zhou, M. (2021b). InfoXLM: An Information-Theoretic Framework for Cross-Lingual Language Model Pre-Training. In Proceedings of the 2021 Conference of the North American Chapter of the Association for Computational Linguistics: Human Language Technologies (pp. 3576–3588).

Clifton, C., Ferreira, F., Henderson, J. M., Inhoff, A. W., Liversedge, S. P., Reichle, E. D., & Schotter, E. R. (2016). Eye movements in reading and information processing: Keith Rayner’s 40-year legacy. Journal of Memory and Language, 86, 1–19.

Collins, C., & Kayne, R. (2011). Syntactic structures of the world’s languages.

Conneau, A., Khandelwal, K., Goyal, N., Chaudhary, V., Wenzek, G., Guzmán, F., Grave, É., Ott, M., Zettlemoyer, L., & Stoyanov, V. (2020). Unsupervised cross-lingual representation learning at scale. Proceedings of the 58th Annual Meeting of the Association for Computational Linguistics, 8440–8451.

Conwell, C., McMahon, E., Vinken, K., Prince, J. S., Alvarez, G., Konkle, T., … & Livingstone, M. (2024). Is the visual cortex really “language-aligned”? Perspectives from Model-to-Brain Comparisons in Human and Monkeys on the Natural Scenes Dataset. Journal of Vision, 24(10), 1288–1288.

Correia, J. M., Formisano, E., Valente, G., Hausfeld, L., Jansma, B., & Bonte, M. (2014). Brain-based translation: fMRI decoding of spoken words in bilinguals reveals language-independent semantic representations in anterior temporal lobe. Journal of Neuroscience, 34(1), 332– 338.

Correia, J. M., Jansma, B., Hausfeld, L., Kikkert, S., & Bonte, M. (2015). Eeg decoding of spoken words in bilingual listeners: From words to language invariant semantic-conceptual representations. Frontiers in psychology, 6, 71.

NLLB Team. (2024). Scaling neural machine translation to 200 languages. Nature, 630(8018), 841.

de Varda, A. G., Berzak, Y., Fedorenko, E., & Levy, R. (in preparation). A Comparison between Surprisal and Language Model Embeddings as Predictors of Behavioral and Neural Responses during Language Processing.

de Varda, A. G., & Marelli, M. (2023). Data-driven cross-lingual syntax: An agreement study with massively multilingual models. Computational Linguistics, 49(2), 261–299.

de Varda, A. G., & Marelli, M. (2024). The Emergence of Semantic Units in Massively Multilingual Models. In Proceedings of the 2024 Joint International Conference on Computational Linguistics, Language Resources and Evaluation (LREC-COLING 2024) (pp. 15910–15921).

de Varda, A. G., Petilli, M., & Marelli, M. (2025). A distributional model of concepts grounded in the spatial organization of objects. Journal of Memory and Language, 142, 104624.

Dehghani, M., Boghrati, R., Man, K., Hoover, J., Gimbel, S. I., Vaswani, A., Zevin, J. D., Immordino-Yang, M. H., Gordon, A. S., Damasio, A., et al. (2017). Decoding the neural representation of story meanings across languages. Human brain mapping, 38(12), 6096–6106.

Deniz, F., Nunez-Elizalde, A. O., Huth, A. G., & Gallant, J. L. (2019). The representation of semantic information across human cerebral cortex during listening versus reading is invariant to stimulus modality. Journal of Neuroscience, 39(39), 7722–7736.

Devlin, J., Chang, M.-W., Lee, K., & Toutanova, K. (2019). BERT: Pre-training of deep bidirectional transformers for language understanding. Proceedings of NAACL-HLT, 4171–4186.

Doerig, A., Kietzmann, T. C., Allen, E., Wu, Y., Naselaris, T., Kay, K., & Charest, I. (2022). Semantic scene descriptions as an objective of human vision. arXiv preprint arXiv:2209.11737, 10.

Duncan, J. (2010). The multiple-demand (MD) system of the primate brain: Mental programs for intelligent behaviour. Trends in Cognitive Sciences, 14(4), 172–179.

Duncan, J., Assem, M., & Shashidhara, S. (2020). Integrated intelligence from distributed brain activity. Trends in Cognitive Sciences, 24(10), 838–852.

Evans, N., & Levinson, S. C. (2009). The myth of language universals: Language diversity and its importance for cognitive science. Behavioral and Brain Sciences, 32(5), 429–448.

Everett, D. (2005). Cultural constraints on grammar and cognition in Pirahã: Another look at the design features of human language. Current anthropology, 46(4), 621–646.

Fedorenko, E., & Blank, I. A. (2020). Broca’s area is not a natural kind. Trends in cognitive sciences, 24(4), 270–284.

Fedorenko, E., Hsieh, P. J., Nieto-Castañón, A., Whitfield-Gabrieli, S., & Kanwisher, N. (2010). New method for fMRI investigations of language: defining ROIs functionally in individual subjects. Journal of neurophysiology, 104(2), 1177–1194.

Fedorenko, E., Duncan, J., & Kanwisher, N. (2013). Broad domain generality in focal regions of frontal and parietal cortex. Proceedings of the National Academy of Sciences, 110(41), 16616–16621.

Fedorenko, E., Ivanova, A. A., & Regev, T. I. (2024a). The language network as a natural kind within the broader landscape of the human brain. Nature Reviews Neuroscience, 1–24.

Fedorenko, E., Piantadosi, S. T., & Gibson, E. A. (2024b). Language is primarily a tool for communication rather than thought. Nature, 630(8017), 575–586.

Fedorenko, E., & Shain, C. (2021). Similarity of computations across domains does not imply shared implementation: the case of language comprehension. Current Directions in Psychological Science, 30(6), 526–534.

Feghhi, E., Hadidi, N., Song, B., Blank, I. A., & Kao, J. C. (2024). What Are Large Language Models Mapping to in the Brain? A Case Against Over-Reliance on Brain Scores. arXiv preprint arXiv:2406.01538.

Fisher, R. A. (1921). On the “probable error” of a coefficient of correlation deduced from a small sample. Metron, 1, 3–32.

Forgey, M. (2024). Quantifying the lack of cross-linguistic diversity in fMRI research (Master’s thesis). Universitat Pompeu Fabra, Barcelona, Spain.

Frank, M. C., Everett, D. L., Fedorenko, E., & Gibson, E. (2008). Number as a cognitive technology: Evidence from Pirahã language and cognition. Cognition, 108(3), 819–824.

Futrell, R., Gibson, E., Tily, H. J., Blank, I., Vishnevetsky, A., Piantadosi, S., & Fedorenko, E. (2018). The Natural Stories Corpus. In Proceedings of the Eleventh International Conference on Language Resources and Evaluation (LREC 2018).

Futrell, R., Mahowald, K., & Gibson, E. (2015). Large-scale evidence of dependency length minimization in 37 languages. Proceedings of the National Academy of Sciences, 112(33), 10336–10341.

Gauthier, J., & Levy, R. (2019). Linking artificial and human neural representations of language. In Proceedings of the 2019 Conference on Empirical Methods in Natural Language Processing and the 9th International Joint Conference on Natural Language Processing (EMNLP-IJCNLP) (pp. 529–539).

Gibson, E., Futrell, R., Jara-Ettinger, J., Mahowald, K., Bergen, L., Ratnasingam, S., … & Conway, B. R. (2017). Color naming across languages reflects color use. Proceedings of the National Academy of Sciences, 114(40), 10785–10790.

Gibson, E., Futrell, R., Piantadosi, S. P., Dautriche, I., Mahowald, K., Bergen, L., & Levy, R. (2019). How efficiency shapes human language. Trends in cognitive sciences, 23(5), 389–407.

Gildea, D., & Temperley, D. (2010). Do grammars minimize dependency length?. Cognitive Science, 34(2), 286–310.

Goldstein, A., Zada, Z., Buchnik, E., Schain, M., Price, A., Aubrey, B., … & Hasson, U. (2022). Shared computational principles for language processing in humans and deep language models. Nature neuroscience, 25(3), 369–380.

Günther, F., Petilli, M. A., Vergallito, A., & Marelli, M. (2020). Images of the unseen: Extrapolating visual representations for abstract and concrete words in a data-driven computational model. Psychological Research, 1–21.

Haspelmath, M., Dryer, M. S., Gil, D., & Comrie, B. (2005). The world atlas of language structures. OUP Oxford.

Hasson, U., Nir, Y., Levy, I., Fuhrmann, G., & Malach, R. (2004). Intersubject synchronization of cortical activity during natural vision. Science, 303(5664), 1634–1640.

Heilbron, M., Armeni, K., Schoffelen, J. M., Hagoort, P., & De Lange, F. P. (2022). A hierarchy of linguistic predictions during natural language comprehension. Proceedings of the National Academy of Sciences, 119(32), e2201968119.

Hirano, Y., Stefanovic, B., & Silva, A. C. (2011). Spatiotemporal evolution of the functional magnetic resonance imaging response to ultrashort stimuli. Journal of Neuroscience, 31(4), 1440–1447.

He, P., Gao, J., & Chen, W. (2022). DeBERTaV3: Improving DeBERTa using ELECTRA-Style Pre-Training with Gradient-Disentangled Embedding Sharing. In The Eleventh International Conference on Learning Representations.

Honey, C. J., Thompson, C. R., Lerner, Y., & Hasson, U. (2012). Not lost in translation: neural responses shared across languages. Journal of Neuroscience, 32(44), 15277–15283.

Honnibal, M., Montani, I., Van Landeghem, S., & Boyd, A. (2020). SpaCy: Industrial-strength Natural Language Processing in Python. 10.5281/zenodo.1212303

Hosseini, E. A., Schrimpf, M., Zhang, Y., Bowman, S., Zaslavsky, N., & Fedorenko, E. (2024). Artificial neural network language models predict human brain responses to language even after a developmentally realistic amount of training. Neurobiology of Language, 1–21.

Huh, M., Cheung, B., Wang, T., & Isola, P. (2024). The platonic representation hypothesis. arXiv preprint arXiv:2405.07987.

Huth, A. G., De Heer, W. A., Griffiths, T. L., Theunissen, F. E., & Gallant, J. L. (2016). Natural speech reveals the semantic maps that tile human cerebral cortex. Nature, 532(7600), 453–458.

Illes, J., Francis, W. S., Desmond, J. E., Gabrieli, J. D., Glover, G. H., Poldrack, R., … & Wagner, A. D. (1999). Convergent cortical representation of semantic processing in bilinguals. Brain and language, 70(3), 347–363.

Ivanova, A. A., Kauf, C., Gao, R., She, J. S., Kean, H. H., Goldhaber, T., … & Fedorenko, E. (2025). Semantic reasoning takes place largely outside the language network. bioRxiv, 2025-12.

Ivanova, A. A., Mineroff, Z., Zimmerer, V., Kanwisher, N., Varley, R., & Fedorenko, E. (2021). The language network is recruited but not required for nonverbal event semantics. Neurobiology of Language, 2(2), 176–201.

Jawahar, G., Sagot, B., & Seddah, D. (2019). What does BERT learn about the structure of language?. In ACL 2019 - 57th Annual Meeting of the Association for Computational Linguistics.

Jeong, H., Sugiura, M., Sassa, Y., Haji, T., Usui, N., Taira, M., Horie, K., Sato, S., & Kawashima, R. (2007). Effect of syntactic similarity on cortical activation during second language processing: A comparison of English and Japanese among native Korean trilinguals. Human Brain Mapping, 28(3), 194–204.

Johns, B. T., & Jones, M. N. (2012). Perceptual inference through global lexical similarity. Topics in Cognitive Science, 4(1), 103–120

Joshi, P., Santy, S., Budhiraja, A., Bali, K., & Choudhury, M. (2020). The State and Fate of Linguistic Diversity and Inclusion in the NLP World. In Proceedings of the 58th Annual Meeting of the Association for Computational Linguistics (pp. 6282–6293).

Kauf, C., Tuckute, G., Levy, R., Andreas, J., & Fedorenko, E. (2024). Lexical semantic content, not syntactic structure, is the main contributor to ANN-brain similarity of fMRI responses in the language network. Neurobiology of Language, 5(1), 7–42.

Kemp, C., Xu, Y., & Regier, T. (2018). Semantic typology and efficient communication. Annual Review of Linguistics, 4(1), 109–128.

Klein, S., & Tsarfaty, R. (2020). Getting the## life out of living: How adequate are word-pieces for modelling complex morphology?. In Proceedings of the 17th SIGMORPHON workshop on computational research in phonetics, phonology, and morphology (pp. 204–209).

Kriegeskorte, N. (2015). Deep neural networks: a new framework for modeling biological vision and brain information processing. Annual review of vision science, 1(1), 417–446.

Kuperman, V., Schroeder, S., & Gnetov, D. (2024). Word length and frequency effects on text reading are highly similar in 12 alphabetic languages. Journal of Memory and Language, 135, 104497.

La Tour, T. D., Eickenberg, M., Nunez-Elizalde, A. O., & Gallant, J. L. (2022). Feature-space selection with banded ridge regression. NeuroImage, 264, 119728.

Lambon Ralph, M. A., Jefferies, E., Patterson, K., & Rogers, T. T. (2017). The neural and computational bases of semantic cognition. Nature reviews neuroscience, 18(1), 42–55.

LeBel, A., Jain, S., & Huth, A. G. (2021). Voxelwise encoding models show that cerebellar language representations are highly conceptual. Journal of Neuroscience, 41(50), 10341–10355.

Levshina, N., & Moran, S. (2021). Efficiency in human languages: Corpus evidence for universal principles. Linguistics Vanguard, 7(s3), 20200081.

Liang, Y., Xu, K., & Ran, Q. (2024). Shared structure of fundamental human experience revealed by polysemy network of basic vocabularies across languages. Scientific Reports, 14(1), 5877.

Lin, X. V., Mihaylov, T., Artetxe, M., Wang, T., Chen, S., Simig, D., Ott, M., Goyal, N., Bhosale, S., Du, J., et al. (2021). Few-shot learning with multilingual language models. arXiv preprint arXiv:2112.10668.

Littell, P., Mortensen, D. R., Lin, K., Kairis, K., Turner, C., & Levin, L. (2017). Uriel and lang2vec: Representing languages as typological, geographical, and phylogenetic vectors. Proceedings of the 15th Conference of the European Chapter of the Association for Computational Linguistics: Volume 2, Short Papers, 8–14.

Lopopolo, A., Van den Bosch, A., Petersson, K. M., & Willems, R. M. (2021). Distinguishing syntactic operations in the brain: Dependency and phrase-structure parsing. Neurobiology of Language, 2(1), 152–175.

Louradour, J. (2023). Whisper-timestamped.

Louwerse, M. M. (2011). Symbol interdependency in symbolic and embodied cognition. Topics in Cognitive Science, 3(2), 273–302.

Majid, A., Roberts, S. G., Cilissen, L., Emmorey, K., Nicodemus, B., O’grady, L., … & Levinson, S. C. (2018). Differential coding of perception in the world’s languages. Proceedings of the National Academy of Sciences, 115(45), 11369–11376.

Malik-Moraleda, S., Ayyash, D., Gallée, J., Affourtit, J., Hoffmann, M., Mineroff, Z., … & Fedorenko, E. (2022). An investigation across 45 languages and 12 language families reveals a universal language network. Nature Neuroscience, 25(8), 1014–1019.

Malik-Moraleda, S., Taliaferro, M., Shannon, S., Jhingan, N., Swords, S., Peterson, D. J., … & Fedorenko, E. (2025). Constructed languages are processed by the same brain mechanisms as natural languages. Proceedings of the National Academy of Sciences

Millet, J., Caucheteux, C., Boubenec, Y., Gramfort, A., Dunbar, E., Pallier, C., & King, J. R. (2022). Toward a realistic model of speech processing in the brain with self-supervised learning. Advances in Neural Information Processing Systems, 35, 33428–33443.

Mineroff, Z., Blank, I. A., Mahowald, K., & Fedorenko, E. (2018). A robust dissociation among the language, multiple demand, and default mode networks: Evidence from inter-region correlations in effect size. Neuropsychologia, 119, 501–511.

Moran, S., McCloy, D., & Wright, R. (2014). PHOIBLE online.

Nakagi, Y., Matsuyama, T., Koide-Majima, N., Yamaguchi, H., Kubo, R., Nishimoto, S., & Takagi, Y. (2024, November). Unveiling Multi-level and Multi-modal Semantic Representations in the Human Brain using Large Language Models. In Proceedings of the 2024 Conference on Empirical Methods in Natural Language Processing (pp. 20313-20338).

Nieto-Castanon, A. (2020). FMRI minimal preprocessing pipeline. In Handbook of functional connectivity Magnetic Resonance Imaging methods in CONN (pp. 3–16). Hilbert Press. doi:10.56441/hilbertpress.2207.6599

Nieto-Castañón, A., & Fedorenko, E. (2012). Subject-specific functional localizers increase sensitivity and functional resolution of multi-subject analyses. Neuroimage, 63(3), 1646–1669.

Niu, J., Lu, W., & Penn, G. (2022). Does BERT rediscover a classical NLP pipeline?. In Proceedings of the 29th International Conference on Computational Linguistics (pp. 3143-3153).

Paunov, A. M., Blank, I. A., Jouravlev, O., Mineroff, Z., Gallée, J., & Fedorenko, E. (2022). Differential tracking of linguistic vs. mental state content in naturalistic stimuli by language and theory of mind (ToM) brain networks. Neurobiology of Language, 3(3), 413–440.

Pereira, F., Lou, B., Pritchett, B., Ritter, S., Gershman, S. J., Kanwisher, N., … & Fedorenko, E. (2018). Toward a universal decoder of linguistic meaning from brain activation. Nature communications, 9(1), 963.

Piantadosi, S. T., Tily, H., & Gibson, E. (2011). Word lengths are optimized for efficient communication. Proceedings of the National Academy of Sciences, 108(9), 3526–3529.

Prince, J. S., Charest, I., Kurzawski, J. W., Pyles, J. A., Tarr, M. J., & Kay, K. N. (2022). Improving the accuracy of single-trial fMRI response estimates using GLMsingle. Elife, 11, e77599.

Radford, A., Kim, J. W., Xu, T., Brockman, G., McLeavey, C., & Sutskever, I. (2023). Robust speech recognition via large-scale weak supervision. International Conference on Machine Learning, 28492–28518.

Ravfogel, S., Elazar, Y., Gonen, H., Twiton, M., & Goldberg, Y. (2020). Null it out: Guarding protected attributes by iterative nullspace projection. In Proceedings of the 58th annual meeting of the association for computational linguistics (pp. 7237–7256).

Regev, M., Honey, C. J., Simony, E., & Hasson, U. (2013). Selective and invariant neural responses to spoken and written narratives. Journal of Neuroscience, 33(40), 15978–15988.

Roads, B. D., & Love, B. C. (2020). Learning as the unsupervised alignment of conceptual systems. Nature Machine Intelligence, 2(1), 76–82.

Rueckl, J. G., Paz-Alonso, P. M., Molfese, P. J., Kuo, W. J., Bick, A., Frost, S. J., … & Frost, R. (2015). Universal brain signature of proficient reading: Evidence from four contrasting languages. Proceedings of the National Academy of Sciences, 112(50), 15510–15515.

Ryskina, M., Tuckute, G., Fung, A., Malkin, A., & Fedorenko, E. (2025), Language models align with brain regions that represent concepts across modalities. In Proceedings of the Second Conference on Language Modeling.

Saha, S., Li, S., Tuckute, G., Li, Y., Zhang, R. Y., Wehbe, L., … & Khosla, M. (2025). Modeling the language cortex with form-independent and enriched representations of sentence meaning reveals remarkable semantic abstractness. arXiv preprint arXiv:2510.02354.

Sánchez, A., Carreiras, M., & Paz-Alonso, P. M. (2023). Word frequency and reading demands modulate brain activation in the inferior frontal gyrus. Scientific Reports, 13(1), 17217.

Sanh, V., Debut, L., Chaumond, J., & Wolf, T. (2019). Distilbert, a distilled version of bert: Smaller, faster, cheaper and lighter. arXiv preprint arXiv:1910.01108.

Saxe, R., Brett, M., & Kanwisher, N. (2006). Divide and conquer: a defense of functional localizers. NeuroImage, 30(4), 1088–1096.

Schrimpf, M., Blank, I. A., Tuckute, G., Kauf, C., Hosseini, E. A., Kanwisher, N., … & Fedorenko, E. (2021). The neural architecture of language: Integrative modeling converges on predictive processing. Proceedings of the National Academy of Sciences, 118(45), e2105646118.

Shain, C., Blank, I. A., Fedorenko, E., Gibson, E., & Schuler, W. (2022). Robust effects of working memory demand during naturalistic language comprehension in language-selective cortex. Journal of Neuroscience, 42(39), 7412–7430.

Shain, C., Blank, I. A., van Schijndel, M., Schuler, W., & Fedorenko, E. (2020). fMRI reveals language-specific predictive coding during naturalistic sentence comprehension. Neuropsychologia, 138, 107307.

Shain, C., & Fedorenko, E. (2025). A language network in the individualized functional connectomes of over 1,000 human brains doing arbitrary tasks. bioRxiv, 2025-03.

Sheikh, U. A., Carreiras, M., & Soto, D. (2019). Decoding the meaning of unconsciously processed words using fMRI-based MVPA. NeuroImage, 191, 430–440.

Sheikh, U. A., Carreiras, M., & Soto, D. (2021). Neurocognitive mechanisms supporting the generalization of concepts across languages. Neuropsychologia, 153, 107740.

Shliazhko, O., Fenogenova, A., Tikhonova, M., Mikhailov, V., Kozlova, A., & Shavrina, T. (2022). mGPT: Few-Shot Learners Go Multilingual. arXiv e-prints, arXiv-2204.

Silva, A. B., Liu, J. R., Metzger, S. L., Bhaya-Grossman, I., Dougherty, M. E., Seaton, M. P., … & Chang, E. F. (2024). A bilingual speech neuroprosthesis driven by cortical articulatory representations shared between languages. Nature Biomedical Engineering, 1-15.

Small, H., Masson, H. L., Mostofsky, S., & Isik, L. (2024). Vision and language representations in multimodal AI models and human social brain regions during natural movie viewing. In UniReps: 2nd Edition of the Workshop on Unifying Representations in Neural Models.

Speer, R. (2022). rspeer/wordfreq: v3.0 (v3.0.2). Zenodo. 10.5281/zenodo.7199437

Stanczak, K., Ponti, E., Hennigen, L. T., Cotterell, R., & Augenstein, I. (2022). Same Neurons, Different Languages: Probing Morphosyntax in Multilingual Pre-trained Models. In Proceedings of the 2022 Conference of the North American Chapter of the Association for Computational Linguistics: Human Language Technologies (pp. 1589–1598).

Stouffer, S. A., Suchman, E. A., DeVinney, L. C., Star, S. A., & Williams Jr, R. M. (1949). The American soldier: Adjustment during army life. Studies in social psychology in World War II, vol. 1.

Sueoka, Y., Paunov, A., Tanner, A., Blank, I. A., Ivanova, A., & Fedorenko, E. (2024). The Language Network Reliably “Tracks” Naturalistic Meaningful Nonverbal Stimuli. Neurobiology of Language, 5(2), 385–408.

Sun, J., Zhang, X., & Moens, M. F. (2023). Tuning In to Neural Encoding: Linking Human Brain and Artificial Supervised Representations of Language. arXiv preprint arXiv:2310.04460.

Tang, J., Du, M., Vo, V., Lal, V., & Huth, A. (2024). Brain encoding models based on multimodal transformers can transfer across language and vision. Advances in Neural Information Processing Systems, 36.

Tenney, I., Das, D., & Pavlick, E. (2019). BERT rediscovers the classical NLP pipeline. In Proceedings of the 57th annual meeting of the association for computational linguistics (pp. 4593–4601).

Tuckute, G., Sathe, A., Srikant, S., Taliaferro, M., Wang, M., Schrimpf, M., … & Fedorenko, E. (2024a). Driving and suppressing the human language network using large language models. Nature Human Behaviour, 8(3), 544–561.

Tuckute, G., Kanwisher, N., & Fedorenko, E. (2024b). Language in brains, minds, and machines. Annual Review of Neuroscience, 47.

Vagharchakian, L., Dehaene-Lambertz, G., Pallier, C., & Dehaene, S. (2012). A temporal bottleneck in the language comprehension network. Journal of Neuroscience, 32(26), 9089–9102.

Van de Putte, E., De Baene, W., Brass, M., & Duyck, W. (2017). Neural overlap of l1 and l2 semantic representations in speech: A decoding approach. NeuroImage, 162, 106–116.

Van de Putte, E., De Baene, W., Price, C. J., & Duyck, W. (2018). Neural overlap of l1 and l2 semantic representations across visual and auditory modalities: A decoding approach. Neuropsychologia, 113, 68–77.

Wang, W., Wei, F., Dong, L., Bao, H., Yang, N., & Zhou, M. (2020). MiniLM: Deep self-attention distillation for task-agnostic compression of pre-trained transformers. Advances in Neural Information Processing Systems, 33, 5776–5788.

Wehbe, L., Blank, I. A., Shain, C., Futrell, R., Levy, R., von der Malsburg, T., … & Fedorenko, E. (2021). Incremental language comprehension difficulty predicts activity in the language network but not the multiple demand network. Cerebral Cortex, 31(9), 4006–4023.

Whitfield-Gabrieli, S., & Nieto-Castanon, A. (2012). Conn: a functional connectivity toolbox for correlated and anticorrelated brain networks. Brain connectivity, 2(3), 125–141.

Wolf, T., Debut, L., Sanh, V., Chaumond, J., Delangue, C., Moi, A., Cistac, P., Rault, T., Louf, R., Funtowicz, M., Davison, J., Shleifer, S., von Platen, P., Ma, C., Jernite, Y., Plu, J., Xu, C., Le Scao, T., Gugger, S., & Rush, A. (2020). Transformers: State-of-the-art natural language processing. In Q. Liu & D. Schlangen (Eds.), Proceedings of the 2020 conference on empirical methods in natural language processing: System demonstrations (pp. 38–45). Association for Computational Linguistics.

Xu, M., Li, D., & Li, P. (2021). Brain decoding in multiple languages: Can cross-language brain decoding work? Brain and Language, 215, 104922.

Xu, X., Wu, C., Rosenman, S., Lal, V., Che, W., & Duan, N. (2023). Bridgetower: Building bridges between encoders in vision-language representation learning. In Proceedings of the AAAI Conference on Artificial Intelligence (Vol. 37, No. 9, pp. 10637–10647).

Xue, L., Constant, N., Roberts, A., Kale, M., Al-Rfou, R., Siddhant, A., Barua, A., & Raffel, C. (2021). MT5: A massively multilingual pre-trained text-to-text transformer. Proceedings of the 2021 Conference of the North American Chapter of the Association for Computational Linguistics: Human Language Technologies, 483–498.

Yamashita, M., Kubo, R., & Nishimoto, S. (2023). Cortical representations of languages during natural dialogue. bioRxiv, 2023-08.

Yang, Y., Wang, J., Bailer, C., Cherkassky, V., & Just, M. A. (2017a). Commonalities and differences in the neural representations of English, Portuguese, and Mandarin sentences: When knowledge of the brain-language mappings for two languages is better than one. Brain and language, 175, 77–85.

Yang, Y., Wang, J., Bailer, C., Cherkassky, V., & Just, M. A. (2017b). Commonality of neural representations of sentences across languages: Predicting brain activation during Portuguese sentence comprehension using an English-based model of brain function. NeuroImage, 146, 658–666.

Youn, H., Sutton, L., Smith, E., Moore, C., Wilkins, J. F., Maddieson, I., … & Bhattacharya, T. (2016). On the universal structure of human lexical semantics. Proceedings of the National Academy of Sciences, 113(7), 1766–1771.

Zinszer, B. D., Anderson, A. J., Kang, O., Wheatley, T., & Raizada, R. D. (2016). Semantic structural alignment of neural representational spaces enables translation between English and Chinese words. Journal of Cognitive Neuroscience, 28(11), 1749–1759.

Zipf, George K. (1949). Human Behavior and the Principle of Least Effort. Cambridge, MA: Addison-Wesley

