## Supplementary information for "Multilingual Computational Models Capture a Shared Meaning Component in Brain Responses across 21 Languages"

#### S1. Related work

A few lines of past research bear relevance to the current work. For example, one approach to evaluating the similarity of representations across languages has been to train a decoder model on brain activity in response to one language, and then to apply this model to brain responses to a different language (see Xu et al., 2021 for a review). For example, Buchweitz et al. (2012) employed Multivoxel Pattern Analysis (MVPA; Haxby et al., 2001) to develop a decoder model based on English and then assessed its effectiveness on translation-equivalent words in Portuguese, reporting an above-chance performance (see also Correia et al., 2014, 2015; Sheikh et al., 2019, 2021; Zinszer et al., 2016). Another approach had relied on inter-subject correlations during naturalistic language comprehension (Hasson et al., 2004). For example, Honey et al. (2012) demonstrated that fMRI time series of English and Russian speakers in response to the same narrative (presented in the two languages) were similar across languages in several temporal, parietal, and frontal cortical areas. These sets of studies converge in showing that the language-processing brain areas respond in a sufficiently similar way to different languages to support generalization across linguistic boundaries.

Focusing on higher-level discourse representations, Dehghani et al. (2017) showed that vector representations of narratives (i.e., embeddings reflecting their high-level meaning) can predict brain responses in the default mode network, and that the story representations supporting the mapping could be switched across languages; however, differently from our procedure, the (decoding) models were not transferred across languages but rather re-trained with story representations from other languages, given that the story representations could not be projected onto the same space. This study therefore leaves open the question of whether the general mapping between linguistic and neural representations is actually shared across languages. Most similarly to our approach, Chen et al. (2024) showed that voxelwise encoding models based on contextual (mBERT) and non-contextual (fastText) word embeddings can be transferred across languages in Chinese-English bilinguals.

These previous studies provide valuable insights into how the brain represents information in different languages, but they suffer from some limitations. First, most past studies have used single words presented in isolation (Buchweitz et al., 2012; Correia et al., 2014, 2015; Sheikh et al., 2019, 2021; Van de Putte et al., 2018; Zinszer et al., 2016), and only a few experiments have used sentence stimuli (Chen et al., 2024; Dehghani et al., 2017; Honey et al., 2012; Yang et al., 2017a, 2017b)—the preferred stimulus of the language network, which responds only weakly to isolated words (e.g., see Fedorenko et al., 2024a for a review). Moreover, most studies have used bilingual speakers and presented the same individuals with linguistic stimuli in both of their languages (Buchweitz et al., 2012; Chen et al., 2024; Correia et al., 2014, 2015; Sheikh et al., 2019, 2021; Van de Putte et al., 2017, 2018; but see Dehghani et al., 2017; Honey et al., 2012; Yang et al., 2017a, 2017b; Zinszer et al., 2016). Especially for single-word stimuli, successful cross-lingual decoding might simply reflect the association of translation equivalents in bilingual speakers, as opposed to genuine similarity in neural representations across languages (see Xu et al., 2021 for a discussion of this limitation). Finally, prior research predominantly utilized pattern classification methods (e.g., MVPA), which are powerful techniques to detect representational regularities across languages but cannot generate predictions for novel stimuli beyond those presented during the training procedure (a similar limitation characterizes the inter-subject correlation approach), whereas encoding models based on contextualized word embeddings can predict brain responses to arbitrary stimuli. Besides these limitations, past studies have at most considered three languages (Dehghani et al., 2017; Yang et al., 2017a). Although the number of languages included in our current study (21 languages across 7 language families) is still only a tiny fraction of the world's ~7,000 languages, we come a little closer to sampling the rich variety that characterizes the human languages.

### S2. Results in individual languages (Study I)

In the main text, we aggregated the encoding results across languages to emphasize the cross-lingual consistencies in encoding performance. This approach was driven by the primary objective of our study, which was to uncover language-general patterns and trends that hold across a variety of languages, and by the sheer volume of the dataset (20 language models, each comprising several layers ranging from 6 to 48, and data spanning 12 languages). This section presents a more granular view of our results. Here, we report the best-layer encoding performance for each language and model combination, as well as the results averaging across language models, both for the WITHIN (**Supplementary Figure 1A**, left) and the ACROSS conditions (**Supplementary Figure 1A**, right).

### S3. Results in individual languages (Study II)

As for Study I, we report the transfer performance for each language and model combination, both for the encoding models trained on passage/story data (**Supplementary Figure 1B**, left) and on sentence data (**Supplementary Figure 1B**, right). Encoding performance is evaluated considering a single layer for each model, i.e., the one that was associated with the strongest encoding performance in Study I (ACROSS condition). Differently from Study I, where mGPT and XGLM were not evaluated in all languages (because some of them were not represented in their pre-training data), in Study II all models were evaluated in all the languages considered.

### S4. WITHIN encoding performance using data from Study II

To evaluate whether language representations from MNNLMs could predict brain responses in the 9 additional languages we considered for Study II, we replicated the WITHIN encoding analysis (Results, "Study I – Language models predict brain responses in diverse languages" and Methods, "Study I – WITHIN encoding models") using the newly collected fMRI data. We kept all the details concerning model fitting (e.g., penalty estimation, normalization, cross-validation approach) identical to Study I with one exception: instead of evaluating all layers for each model, we only considered a single layer per model, namely the layer which obtained the best performance in Study I. Since in Study II we collected brain responses to three passages in each language (some of which were discarded due to low data quality, see

Methods, “Study II – Time series extraction”), we averaged encoding performance scores across passages, excluding those that did not pass the selection criteria.

Because Study II considers responses between two and three participants per language (depending on correlations in the time series; see Methods, “Study II – Time series extraction”), we adapted the cross-participant generalization scheme. For each passage and language, we performed leave-one-participant-out (LOO) splits that train on 90% of the data from two participants and test on 10% of the data from the third participant (or train on one and test on one when only two participants passed quality control). For each language, we averaged across that language’s kept passages (between 1 and 3; see Methods, “Study II – Time series extraction”). Finally, for each model we averaged the encoding scores across languages. Statistical significance was assessed against four circular-shift baseline models as in the main analyses.

Confirming our observations from Study I, our results showed that the MNNLMs we considered were able to predict fMRI responses in most languages (**Supplementary Figure 3A**). Aggregating the results across languages, the majority of MNNLMs were significantly predictive of brain responses (all $p < 0.05$ apart from DistilmBERT and $mT5_{base}$; **Supplementary Figure 3B**). Once again, large auto-regressive models obtained strong predictive performance (mGPT: $r = 0.19$, $SE = 0.04$, $p < 0.001$; $XGLM_{xl}$: $r = 0.19$, $SE = 0.04$). The encoding performance obtained by the various models on the Study II data was significantly correlated with the encoding scores obtained in Study I ($r = 0.79$, $p < 0.001$, **Supplementary Figure 3C**), indicating that inter-model differences in brain predictivity are highly reliable.

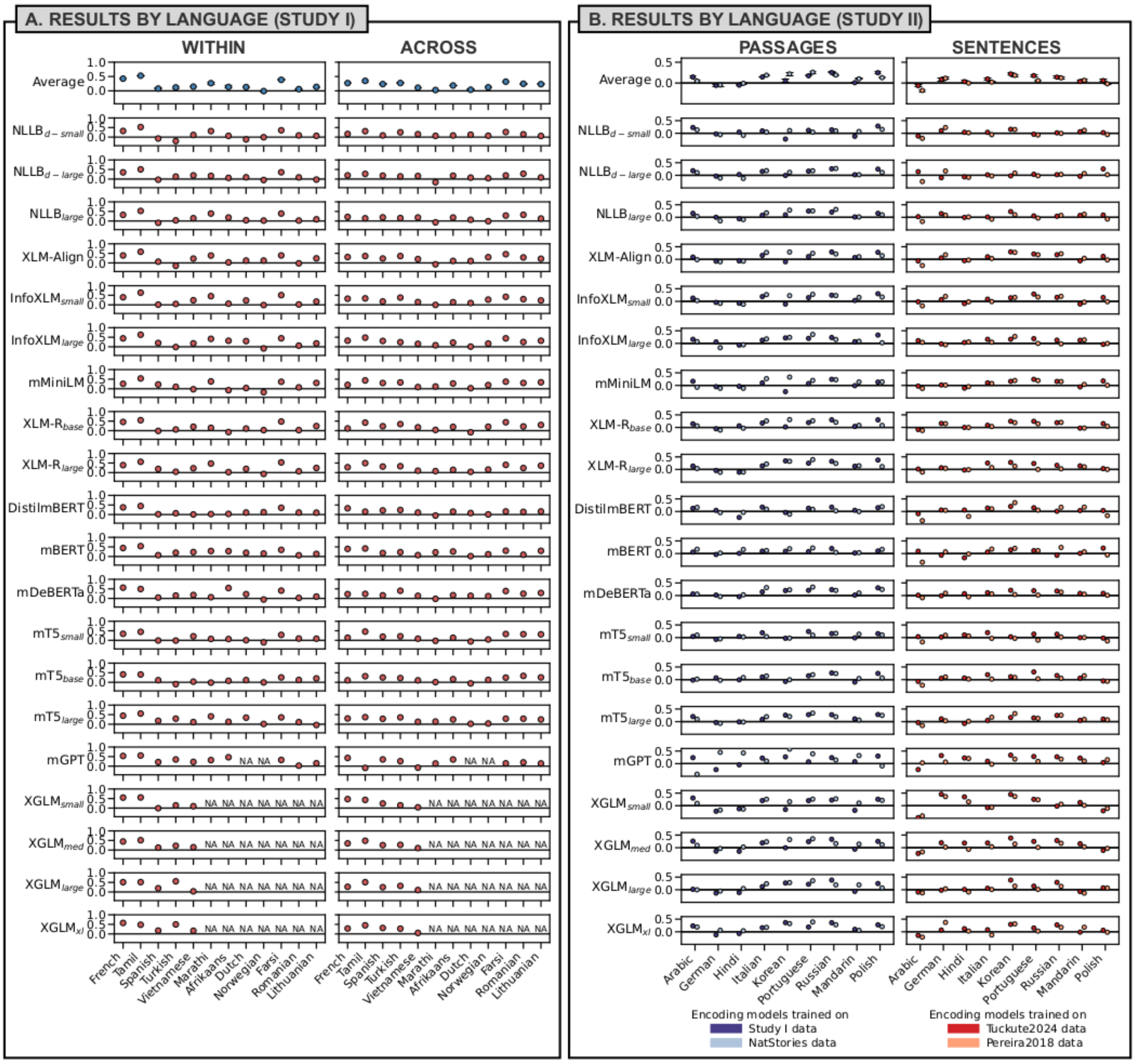


**Supplementary Figure 1.** Encoding results by language. **A.** Best-layer encoding performance scores for each model and language, for the WITHIN (left) and ACROSS (right) condition. The rows on the top report the language-wise encoding performance averaged across models, with error bars indicating the standard error of the mean across models. Note that XGLM encoding scores are not available for Marathi, Afrikaans, Dutch, Norwegian, Farsi, Romanian, and Lithuanian, and mGPT encoding results are not available for Dutch and Norwegian. **B.** Transfer performance of the encoding models trained on passages (left; *Study I*: dark blue, *NatStories*: light blue) and sentences (right; *Tuckute2024*: dark red; *Pereira2018*: pink) obtained in each individual language considered in Study II. As in panel A, the rows on the top report the language-wise encoding performance averaged across models, with error bars indicating the standard error of the mean across models.

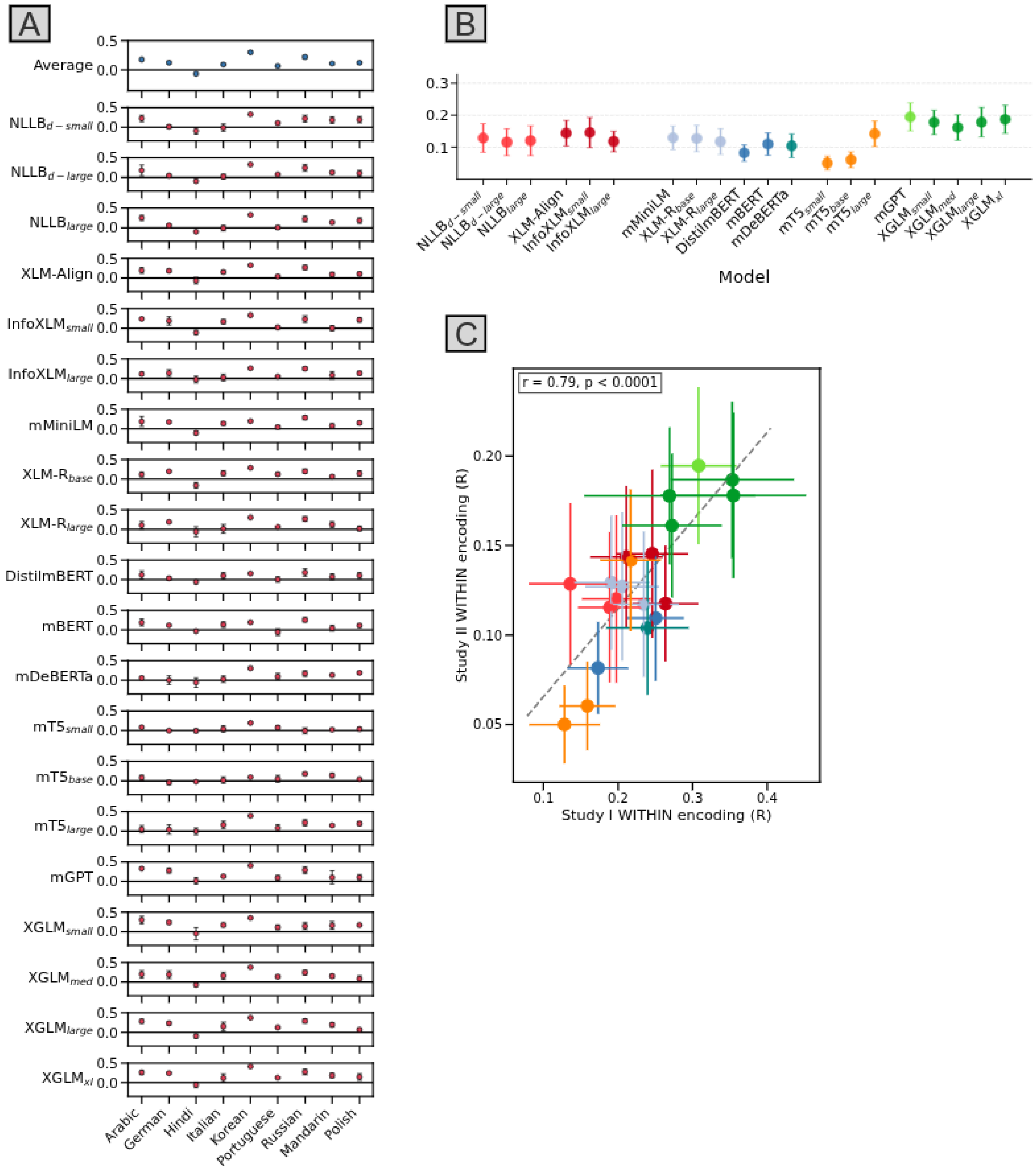


**Supplementary Figure 3:** WITHIN encoding performance obtained on the fMRI data from Study II. *A*: Encoding scores for each model and language, averaged across passages. The error bars indicate the standard error of the mean of the encoding performance across passages. The row on top reports the encoding performance averaged across models, with the error bars indicating the standard error of the mean across models. *B:* Model-wise summary of the encoding results. *C:* Relationship between each model's WITHIN encoding performance obtained on the data from Study I ($x$-axis) and Study II ($y$-axis). The error bars report the standard error of the mean of the encoding performance along the two axes.

### S5. Performance in the median layer

In the main body of our paper, we presented the results obtained by the best-performing layer of each neural language model. This approach highlights the peak capabilities of each model, but it is also important to consider some indices of the central tendency of the models' performance. To this end, we report the results of the median-performing layer of each model. This analysis is relevant given that models have a varying number of layers, and larger models with more layers could potentially show a wider range of performance levels (and thus, higher *maxima*). Focusing on the median layer mitigates the impact of this variability.

The encoding performance scores for the median layers of each model across individual fROIs are detailed in **Supplementary Figure 4**. Consistent with the best-layer results, median-layer performance shows robust encoding across all five left-hemisphere language regions. In the WITHIN condition, nearly all models achieve significant encoding performance across individual fROIs, with the strongest effects once again in the posterior temporal cortex (average $r$ = 0.19, with 19/20 models significant; **Supplementary Figure 4**, top) and middle frontal gyrus (average r = 0.18, with all 20 models significant). The ACROSS condition shows similarly robust transfer, with all models achieving significant performance when averaged across the language network (average $r$ = 0.16, all $p < 0.001$; **Supplementary Figure 4**, bottom), and strong regional effects particularly in the posterior temporal cortex (average $r$ = 0.19, all $p < 0.001$). Although median-layer performance is slightly lower than best-layer results, the pattern of regional specificity is preserved, confirming that our main findings are not dependent on the choice of the layers.

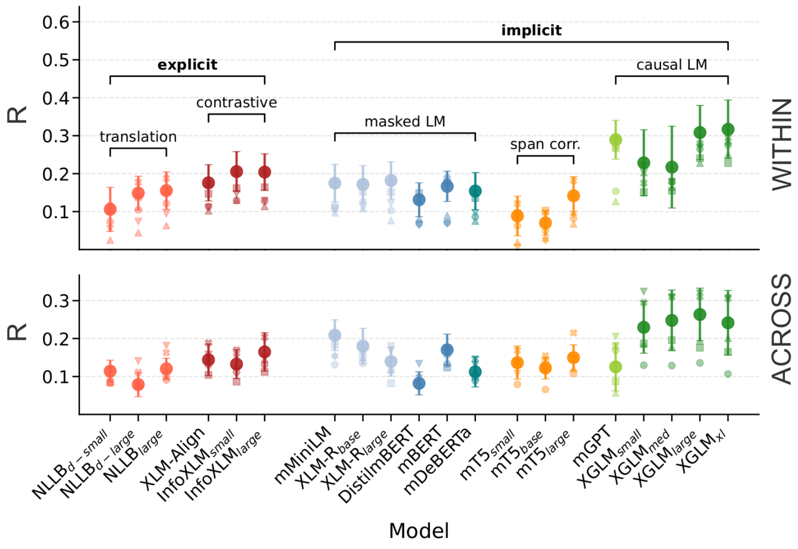


**Supplementary Figure 4.** Encoding results obtained by the median-performing layer in each model under consideration, both in the WITHIN (top) and in the ACROSS (bottom) condition. The scores are averaged across languages. The error bars indicate the standard error of the encoding performance across languages.

### S6. Word frequency, length, and timing baselines

To assess the extent to which brain responses could be captured by surface linguistic and speech properties, we fit simple encoding models where brain activity was predicted by word-level frequency and length, averaged over TRs, as well as word rate and word onset. Word length was defined as the number of characters in each word, whereas frequency (Zipf-transformed) was obtained with the Python package wordfreq (version 3.0), a library for looking up word frequencies in many languages (Speer, 2022). To further control for information tied to the timing of naturalistic audio, we additionally included word rate (number of words per TR) and word onset (onset of the first token in each TR). All other details concerning pre-processing, normalization, model fitting, etc. were left unaltered with respect to the main approach detailed in the Methods ("Study I – Encoding models").

For Study I, brain responses were predicted both within a single language, with sequential 10-fold cross-validation (as in the WITHIN condition), and by transferring the frequency-length models to novel languages (as in the ACROSS condition). Results were averaged across languages. Afrikaans and Marathi were discarded from the analyses because frequency estimates for these two languages were not available through the wordfreq package. In the WITHIN condition, the baseline model achieved $r = 0.14$, $SE = 0.08$ (cf. $r = 0.35$, $SE = 0.08$ obtained by $XGLM_{XL}$ in the same condition and an across-model average of $r = 0.23$) and was at chance in the ACROSS condition ($r = -0.01$, $SE = 0.01$).

For Study II, materials were written in two out of the four sources (*Pereira2018*, *Tuckute2024*); thus, we did not include word onset or word rate, and trained the encoding models based on frequency and word length alone. We trained the encoding models predicting brain responses from those two features on the training data (Study I data, *NatStories*, *Pereira2018*, *Tuckute2024*), and tested them on the newly collected data, following the general procedure we used for the standard encoding analyses performed for Study II (Methods, "Study II – Brain encoding models"). Results were averaged across passages and languages. The models based on frequency and length achieved little to no predictivity across all four training datasets (Study I data: $r = 0.06$, $SE = 0.03$; *NatStories*: $r = -0.01$, $SE = 0.02$; *Pereira2018*: $r = -0.07$, $SE = 0.03$; *Tuckute2024*: $r = 0.01$, $SE = 0.02$).

### S7. Native functional regions of interest

Our main analyses used fROIs defined on the basis of the English localizer to match prior work and ensure comparability across participants. Nevertheless, to address the concern that English-defined fROIs might bias the results toward voxels that respond similarly across languages, we repeated Study I encoding analyses using fROIs defined with the native-language auditory localizer (*intact* > *degraded speech*) from Malik-Moraleda, Ayyash, et al. (2022). For each participant, we defined five language fROIs using the same parcel-constrained, participant-specific procedure as for the English localizer (Fedorenko et al., 2010). We trained encoding models using the identical cross-participant scheme and the same randomized baselines (circular shifts) as the main analyses.

Time series extracted from English- vs. native-defined fROIs were highly correlated across languages: PostTemp $r = 0.78$; AntTemp $r = 0.74$; IFG $r = 0.69$; IFGorb $r = 0.64$; MFG $r = 0.66$; network average $r = 0.79$. These results are in line with prior evidence that English and native localizers pick out similar voxels and responses.

Using these native-language fROIs, we carried out the full set of encoding analyses in Study I. The overall pattern of results was strikingly similar to the one obtained with the English localizer. Model-wise encoding scores were strongly correlated across the two localizers ($r = 0.95$ in the WITHIN condition and $r = 0.83$ in the ACROSS condition, both $p$s $< 0.001$). All of the key findings were preserved: encoding was robust across languages, zero-shot transfer was successful, auto-regressive models outperformed other architectures, larger models outperformed smaller ones, and encoding was strongest in temporal rather than frontal fROIs (**Supplementary Figure 5**). The only notable difference was that encoding scores were slightly lower with the native-language localizer (WITHIN: mean $r$ = 0.18 vs. 0.23; ACROSS: mean $r$ = 0.18 vs. 0.22). We attribute this to lower reliability of the native-language localizer compared to the extensively validated English one, which benefits from decades of testing across many participants.

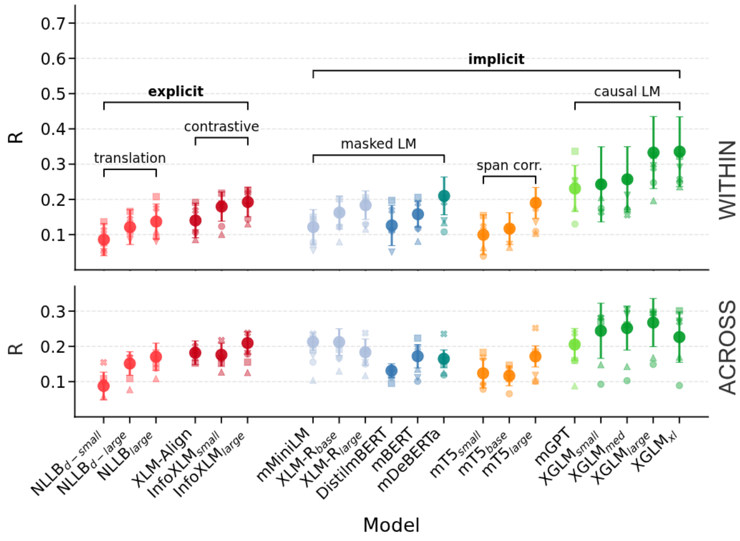


**Supplementary Figure 5.** Encoding results obtained using time series from the fROIs identified in the native language, both in the WITHIN (top) and in the ACROSS (bottom) condition. The scores are averaged across languages. The error bars indicate the standard error of the encoding performance across languages.

### S8. Results within individual fROIs.

In the main body of the paper, we focused on the results obtained in the language areas; here, we report in detail the results obtained in the individual fROIs.

In the WITHIN condition, encoding performance is stronger in the posterior temporal language fROI, reaching an average of $r$ = 0.23 across models (ranging from 0.15 to 0.35, $SE$ = 0.03–0.07; see “Language models predict brain responses in diverse languages”). The anterior temporal fROI yields an average $r$ = 0.19 (ranging from 0.12 to 0.28, $SE$ = 0.04–0.13), with nearly all models significantly above baseline. Responses in the fROIs located in the inferior frontal and middle frontal gyri are also reliably predicted by the encoding models, though at slightly lower levels (both averaged $r$ = 0.17; IFG fROI range: 0.10–0.30, $SE$ = 0.04–0.09; MFG fROI range: 0.09–0.33, $SE$ = 0.04–0.10), with most models significant. The fROI in the orbital part of the IFG shows the weakest encoding performance, with an average of $r$ = 0.15 (range 0.06–0.29, $SE$ = 0.05–0.11), though several models are still above baseline.

In the ACROSS condition, effects are again strongest in the posterior temporal fROI (average $r$ = 0.25 across models; range: 0.15–0.35; $SE$ = 0.03–0.07), followed by the anterior temporal fROI (mean $r$ = 0.18, range 0.05–0.25, $SE$ = 0.04–0.09). Responses in the frontal regions are reliably predicted but at lower levels overall (mean $r$s = 0.16–0.22, ranges 0.08–0.36, $SE$ = 0.03–0.10). These results confirm that ACROSS transfer is distributed across the fronto-temporal language network, with temporal regions showing the strongest alignment.

In Study II, the overall pattern replicates the findings of Study I: encoding is strongest in temporal regions and consistently reliable across the whole fronto-temporal language network.

When training encoding models on data from Study I, performance is highest in the posterior temporal fROI (average $r = 0.14$ across models, range 0.08–0.24, $SE = 0.02$–0.05), followed by the anterior temporal fROI ($r = 0.08$, 0.03–0.15, $SE = 0.03$–0.06). Frontal areas also showed robust alignment, with the IFG and MFG both around $r = 0.07$–0.08 (ranges 0.02–0.14; $SE = 0.03$–0.07), and the orbital IFG at $r = 0.09$ (0.00–0.15; $SE = 0.03$–0.07). All fROIs were significant across nearly all models.

Concerning encoding models trained on *NatStories*, effects were larger overall, with the posterior temporal region again showing highest performance ($r = 0.17$, 0.08–0.24, $SE = 0.02$–0.08), followed by the anterior temporal fROI ($r = 0.15$, 0.06–0.24, $SE = 0.02$–0.09). The IFG and MFG were slightly weaker ($r \approx$ 0.09–0.10, $SE = 0.03$–0.12), and the orbital IFG reached $r = 0.12$ (0.02–0.26, $SE = 0.03$–0.08). Nearly all models performed significantly above baseline.

The *Tuckute2024* dataset exhibited the highest encoding in the anterior temporal fROI ($r = 0.13$, 0.05–0.21, $SE = 0.03$–0.11) and orbital IFG ($r = 0.14$, 0.09–0.19, $SE = 0.03$–0.07). The posterior temporal region followed with $r = 0.08$ (0.00–0.13, $SE = 0.02$–0.08), and the IFG and MFG showed weaker but consistent effects ($r \approx 0.04$–0.08, $SE = 0.01$–0.10).

In the *Pereira2018* dataset, effect sizes were smaller overall, consistent with the results found at the level of the whole language network (see "Study II"). The IFGorb and IFG yielded the highest predictivity ($r = 0.10$–0.15, $SE = 0.02$–0.06), while the anterior and posterior temporal fROIs showed more modest alignment ($r \approx 0.02$–0.09, $SE = 0.01$–0.07). Nevertheless, most models remained above baseline in these regions, underscoring the robustness of the mapping even under noisier conditions.

Pooling across all datasets, encoding was strongest in the posterior and anterior temporal fROIs (mean *r*s = 0.10 each; ranges 0.04–0.17; *SE*s = 0.03–0.07), with reliable though weaker prediction in the frontal regions (*r*s ≈ 0.05–0.11).

### S9. Optimal TR shift

The hemodynamic response measured with fMRI peaks several seconds after the presentation of a stimulus (e.g., Hirano, Stefanovic, & Silva, 2011). Previous work in computational language neuroscience has addressed the issue by either (a) implementing an arbitrary shift corresponding to 6 seconds (Schrimpf et al., 2021), or (b) concatenating embedding representations of the stimuli corresponding to the current and previous temporal resolution intervals to estimate the hemodynamic response function directly from the embeddings (e.g., Aw & Toneva, 2023). However, the approach in (a) does not ensure that the assumed temporal delay is optimal, and the approach in (b) increases the dimensionality of the predictors, raising possible concerns about over-fitting especially in settings where the amount of data is limited. To overcome these limitations, we have identified in a separate dataset the optimal temporal shift for the specific task at hand, i.e., MNNLM-based encoding of brain responses in a naturalistic listening task. We have conducted these analyses on the *NatStories* fMRI corpus (see Methods, "NatStories"). Our procedure was analogous to what we reported in the Methods section; we averaged the fMRI time series across voxels for each fROI, and subsequently across fROIs within the language network. We additionally averaged the network-level time series across participants. We obtained word-by-word speech transcriptions with Whisper-timestamped, and used XGLMsmall to derive word embeddings for the obtained texts, which were then aligned with the fMRI responses by averaging the embeddings over the TR intervals. As a final step, we fit independent encoding models for each temporal shift to identify the optimal delay. We tested five different delays: 0, 1, 2, 3, and 4 TR intervals, equivalent to 0, 2, 4, 6, and 8 seconds, respectively. For instance, with a delay of 1 TR interval, we used the embeddings corresponding to the $i^{th}$ TR as predictors and the fMRI response corresponding to the $i$+1$^{\text{th}}$ image as a response. Similarly, with a delay of 2 TR intervals, the embeddings for the $i^{th}$ TR were aligned with the fMRI response of the $i$+2$^{\text{th}}$ image, and so forth. This systematic evaluation allowed us to determine the optimal temporal alignment between the neural language model embeddings and the fMRI responses, ensuring that our encoding models were accurately capturing the temporal dynamics of the hemodynamic response. The encoding models were fit with cross-validation, where the linear encoding models were iteratively fit in eight out of nine stories, and evaluated on the held-out story.

Performance of the encoding models increased with the temporal delay up to a shift of 3 TR intervals (see **Supplementary Figure 7**). Specifically, we observed a progressive improvement in the model's ability to predict fMRI responses as the temporal delay increased from 0 to 3 TRs. However, performance decreased with a shift of 4 TR intervals, supporting the notion that the optimal temporal delay for capturing the hemodynamic response in our task is around 6 seconds. Therefore, for the main analyses of this paper, we adopted a temporal shift of 3 TR intervals (6 seconds).

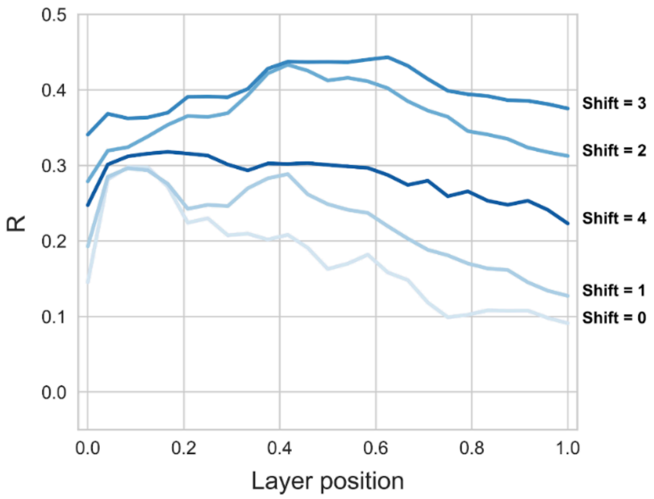


**Supplementary Figure 7.** Layer-wise encoding performance as obtained by XGLMsmall on held-out passages in the *NatStories* fMRI data. We evaluated five different TR shifts to account for the lag in the hemodynamic response, corresponding to 0, 2, … 8 seconds. We found that the optimal encoding performance was obtained with a 3-TR shift, corresponding to a lag of 6 seconds.

### S10. Time series correlation

To pre-select the fMRI data for our analyses, we examined the reliability of BOLD responses to the linguistic stimuli. In particular, we calculated the Pearson correlation between the time series of the two (Study I) or three (Study II) participants in each language (Methods, "Study I – Time series extraction" and "Study II – Time series extraction"). In Study I (where two participants listened to a single passage for each language), the languages where the time series of the two participants did not show a significant correlation were excluded from our sample; in Study II (where three participants listened to three passages for each language), we used this criterion to select participants and passages for each language. As a sanity check, we also compared these *within-language* correlations to correlations computed *across different languages*. Here, we correlated the fMRI time series from one participant with those from participants of other languages (for Study II, we performed this comparison across passages as well, restricting to the same language–passage combinations used in the main analyses). Because the passages were not temporally aligned in different languages, and because all passages were trimmed to the same length (260 s = 130 TRs), the resulting time series are aligned in length but should not be correlated. As expected, the across-language correlations cluster around zero, confirming that the ACROSS encoding results are not trivially explained by correlations in the raw fMRI time series.

**Supplementary Figure 8** summarizes the results for Study I (**Supplementary Figure 8A**) and Study II (**Supplementary Figure 8B**). The correlations within each language (the 'Within' dots) are consistently higher than the average correlation observed between each participant's responses in one language and all other participants across different languages (the 'Avg' dots); the latter correlations cluster around zero.

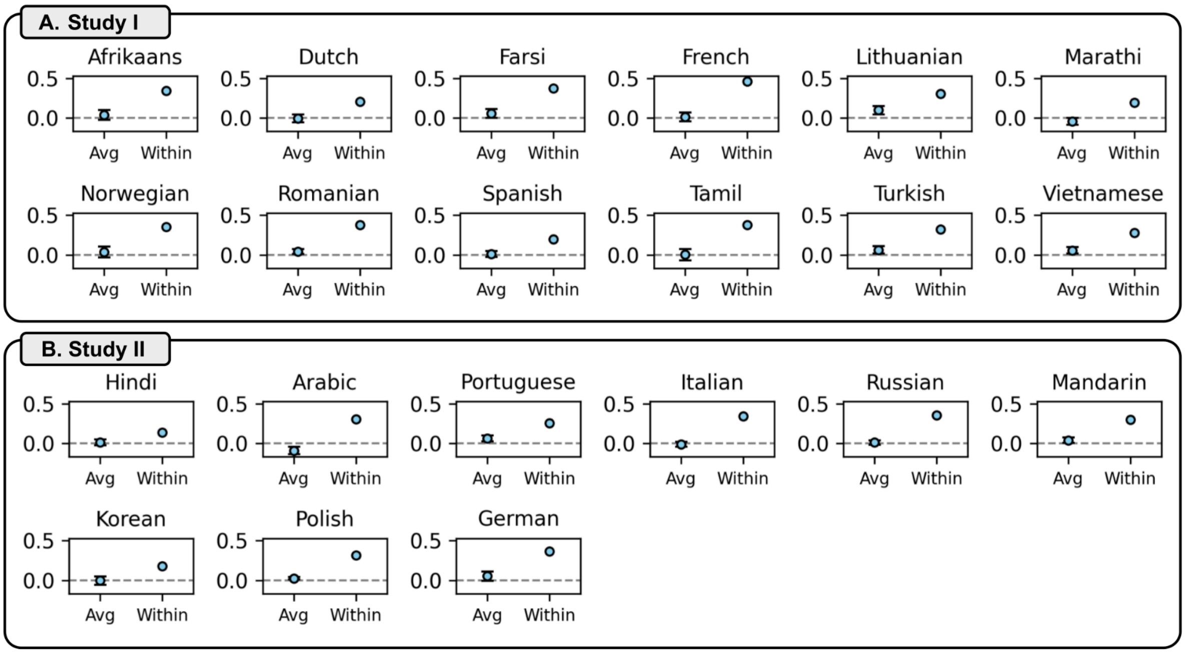


**Supplementary Figure 8.** Comparison of the correlation coefficients between participant pairs within and across languages, both for Study I (A, top) and Study II (B, bottom). The 'Within' dots represent the correlation between the time series of two participants in the same language. The 'Avg' dots show the average correlation between each participant's fMRI responses in one language and those of all other participants across different languages. Error bars indicate the standard error of the mean of these correlations across languages. In the case of Study II (B, bottom), we averaged the correlations across passages. The plot shows that the time series of two participants listening to the same passage are correlated, but the correlation across languages is low.

### S11. Context contamination

A critical aspect often overlooked in computational language neuroscience is the potential for context contamination in transformer-based encoding models. This section presents an approach to evaluate the extent of such contamination.

Context contamination might arise when self-attention mechanisms in language models retain information from previous data folds in a cross-validation setting. If the input text is processed in a single pass, the embeddings generated by the model may carry over contextual information from one fold to the next. This carry-over can confound results, leading to an overestimation of a model's true encoding performance, as it may partially rely on residual information from preceding folds rather than on the current input. Note that, in our approach, context contamination might arise only in the WITHIN setting, as the cross-lingual encoding models are trained and evaluated on different passages in different languages. Nevertheless, we tested both conditions using a more stringent embedding protocol.

To address this potential confound, we refit our encoding models with a modification: resetting the context after each fold when obtaining the models' embeddings. This means that when extracting embeddings for each test fold, the model processes only that fold in isolation (thus preventing carry-over of information from prior tokens). We then re-ran both the WITHIN and ACROSS encoding analyses using these embeddings, keeping all other parameters (including participant-level cross-validation and 10-fold splits) identical to the main analysis. In the case of bidirectional models, a causal setup was simulated as in the main analyses.

It is important to note that, in the context of fMRI data collection, participants were exposed to a continuous stream of language stimuli, allowing for a natural accumulation of context. However, our models, with the reset applied, will not have access to such prior input.

The results obtained by resetting the models' context at the fold boundaries are reported in **Supplementary Figure 9**. The figure displays a contained decline in performance with respect to what we reported in the main analyses (horizontal dotted lines, see also **Figure 2A**).

In the WITHIN condition, the encoding performance averaged across models is $r = 0.13$, compared to $r = 0.23$ obtained with the full context (42% decrease). The posterior temporal fROI remained highest (mean $r = 0.15$), with inferior/middle frontal lower (means $r = 0.10$ and $r = 0.09$). In the ACROSS condition, the reduction in performance was more limited (across-model mean $r = 0.18$ (vs. $r = 0.22$ without reset; 18% decrease). Again, encoding performance was highest in the posterior temporal cortex (mean $r = 0.23$), with frontal regions lower (means $r = 0.11$–$0.15$).

With the available data, it is difficult to determine whether the decrease in encoding performance under context reset reflects context contamination or instead a mismatch between how models and humans process linguistic input (models being limited to constrained windows, humans having access to the full passage). Both WITHIN- and ACROSS-language analyses show lower scores under the reset procedure, which rules out contamination as the sole explanation, since in the ACROSS case train and test passages are entirely different. At the same time, the difference in performance against the full-context baseline is larger in the WITHIN than in the ACROSS condition (42 vs. 18%), and this difference could be due to context contamination proper. Overall, the decline is relatively contained, the ordering of models is preserved, and performance remains reliably above baseline, indicating that neither contamination nor limited context access plays a decisive role in driving the main findings.

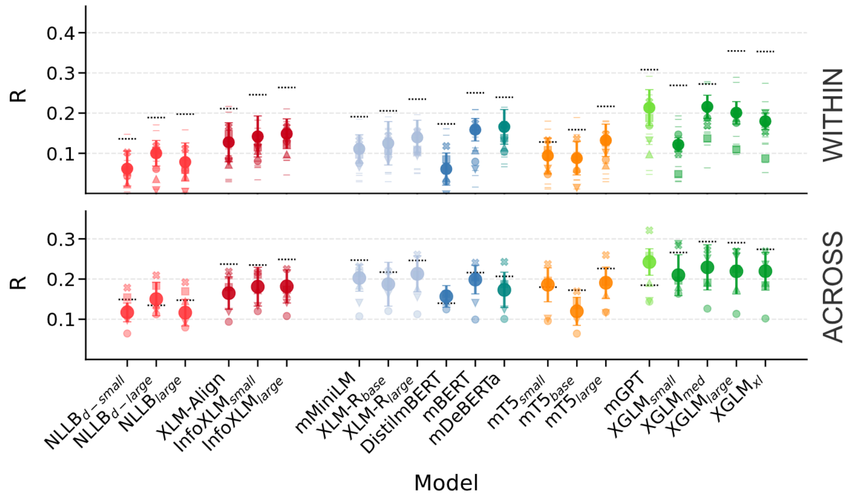


**Supplementary Figure 9.** Best-layer WITHIN (top) and ACROSS (bottom) encoding performance obtained after resetting the models' context across folds. The scores are averaged across languages. The error bars report the standard error of the encoding performance across languages. The horizontal, dotted lines on top of the dots indicate the performance obtained by the various models in the standard setting, without the context being reset at fold boundaries (thus corresponding to the results reported in Figure 2A).

### S12. Data reliability

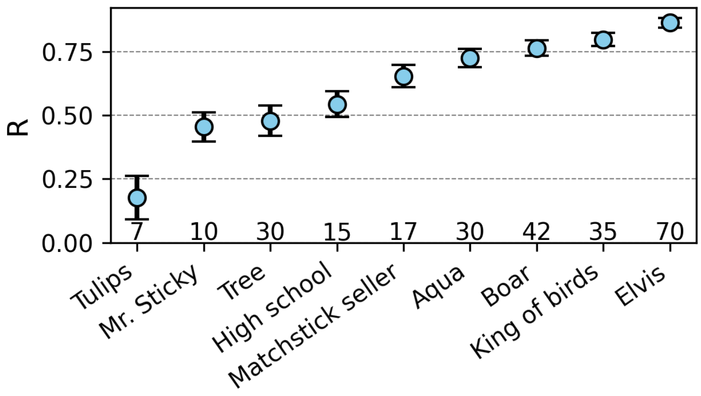


**Supplementary Figure 10.** Split-half reliability for each passage in the *NatStories* fMRI corpus. At the base of each dot, we report the number of participants for each passage.

To get a sense of the quality of the fMRI data employed across Study I and Study II, we conducted a reliability analysis. The results of the analysis are reported in **Figure 1C**.

In the case of Study I, we measured the correlations between the time series of the two participants in each of the 12 languages considered (Methods, "Study I – Time series extraction") and averaged them. For Study II, we adopted the same approach, but for the passages where the time series were reliable for three participants, we averaged the correlations across the three possible participant pairs. The final reliability estimate was obtained by first averaging across participants, then across passages (only considering the passages included in the final sample, see Methods, "Study II – Time series extraction"), and lastly across languages.

For the additional datasets considered for Study II (*Pereira2018*, *Tuckute2024*, and *NatStories*), which included a larger number of participants, we calculated the split-half reliability of the responses. In the case of *NatStories*, for each passage, we randomly assigned the participants to two different groups, averaged their time series within the two groups, and calculated their correlation. This procedure was repeated 1,000 times to obtain more stable estimates. This analysis showed that the majority of the passages obtained a high split-half reliability score (see **Supplementary Figure 10**). An aggregated reliability score was obtained as a weighted average of the reliability scores of each passage, with the weights being proportional to the number of TR-level responses for each passage (ranging from 151 to 213). A similar procedure was adopted for *Pereira2018* and *Tuckute2024*, with the key difference that for these two datasets each datapoint corresponded to a response to a single sentence. *Pereira2018* includes data from two studies (Experiment 2 and Experiment 3), with different numbers of participants (9 and 6). Reliability was estimated independently in each dataset and then aggregated with a weighted average, with the weights being proportional to the number of sentences in each experiment (384 and 243).

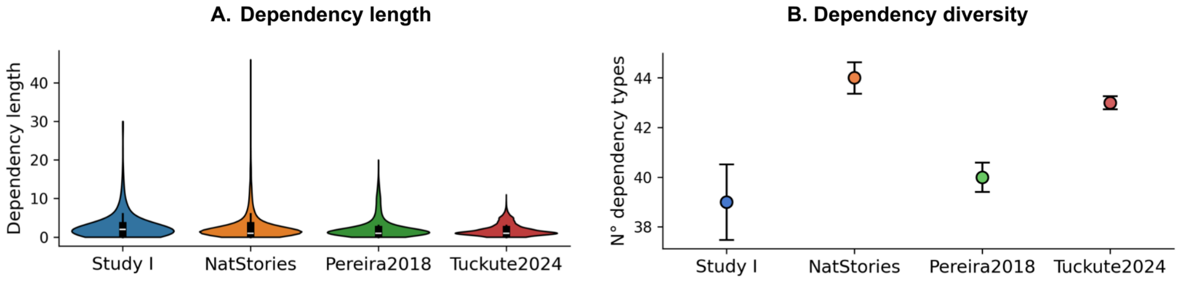


**Supplementary Figure 11.** *A:* Distribution of dependency length across the four datasets employed in Study II. *B:* Number of unique dependency types across the four datasets. The error bars indicate the bootstrapped standard deviation as calculated across 1,000 samples with replacement.

### S13. Syntactic complexity

To measure variation in syntactic complexity across the different datasets employed in Study II (Study I data, *NatStories*, *Pereira2018*, and *Tuckute2024*), we used two complementary metrics: dependency length and dependency diversity. Dependency length measures the linear distance between tokens and their syntactic heads, whereas dependency diversity quantifies the variety of dependency relations within each dataset, reflecting the range of grammatical constructions.

Sentences in all datasets were tokenized and parsed using the SpaCy dependency parser (en_core_web_sm). For each sentence, dependency length was calculated as the absolute difference between the index of a token and its syntactic head. To quantify the diversity of syntactic structures, dependency relations (e.g., *nsubj*, *dobj*, *prep*) were extracted for all tokens in each dataset. The number of unique dependency types in each dataset was calculated as a proxy for syntactic diversity.

This analysis reveals differences in both syntactic depth and diversity across datasets. The *NatStories* dataset includes syntactic dependencies that are both longer, especially in their extremes

(**Supplementary Figure 11A**) and more diverse (**Supplementary Figure 11B**). The Study I dataset comprises syntactic dependencies that are intermediate in length, but with low variability in dependency types. The sentences in the *Pereira2018* and *Tuckute2024* datasets are characterized by short dependencies; furthermore, the *Pereira2018* dataset includes comparatively fewer unique dependency relations.

### S14. Single-language training control and typological factors

In our main analyses (Study I–II), cross-lingual transfer performance was evaluated by training encoding models in all but one language and testing them zero-shot in the held-out language. To assess the robustness of these findings under a more stringent setup, we carried out a control analysis in which encoding models were trained on a single language and transferred to all other languages. This approach represents a stricter test of cross-lingual generalization because the encoding models are estimated from substantially less training data (one language rather than N−1).

For each language *li*, we trained encoding models on 90% of the fMRI data from that language and then tested them zero-shot on the left-out 10% of another language *lj*. This produced two transfer scores for each language pair: *li* → *lj* and *lj* → *li*. Then, we averaged them to obtain a symmetric transferability metric for each pair. Significance was assessed by comparison with circular-shift baselines (see Methods).

**Supplementary Figure 12A** reports the average encoding performance across models, families, and fROIs under this stricter scheme. Network-level performance across models was robust (mean $r$ = 0.113, *SE* = 0.02, all $p < 0.01$), although lower than both the performance obtained in the WITHIN condition (thus using the same cross-validation scheme and amount of data, but testing within a single language; $r$ = 0.23, *SE* = 0.01) and in the ACROSS condition (which similarly involved generalization across languages, but training in N–1 languages; $r$ = 0.22, *SE* = 0.011). Despite the reduction in training data, the overall pattern was highly consistent with our main results: auto-regressive models (XGLM, mGPT) and larger-capacity architectures achieved the strongest cross-lingual performance, followed by other encoder-only and encoder-decoder models. Across fROIs, temporal areas again yielded the strongest alignment, with average correlations up to $r$ = 0.26 for $\text{XGLM}_{\text{med}}$ and $\text{XGLM}_{\text{xl}}$ in the posterior temporal cortex.

**Typological factors.** We next asked whether transfer performance between two languages correlated with several measurements of similarity between the two languages. In particular, we considered:

1. **Genetic distance.** A distance metric based on the Glottolog's language tree. The metric calculates the distance between two languages by counting the number of upward steps required on the tree until both languages converge at a common node. This number is then divided by the total number of branches from the first language to the tree's root.
2. **Geographical distance.** The distance on the earth surface between the geographical locations where two languages are spoken.

3. **Syntactic distance.** This metric measures the pairwise distance between two languages as the cosine distance between feature vectors describing the languages' syntax. These features encode attributes such as word order or the presence of specific constructions. The syntactic features are derived from the World Atlas of Language Structures (WALS; Haspelmath et al., 2005), the Syntactic Structures of the World's Languages (SSWL; Collins & Kayne, 2011), and Ethnologue (Campbell, 2008).
4. **Phonological distance.** Similarly to syntactic distance, phonological distance is computed over phonological feature vectors based on WALS and Ethnologue characterizing the languages.
5. **Phonetic distance.** This distance metric measures the distance between two languages as the cosine distance between the phonetic feature vectors derived from PHOIBLE (Moran, McCloy, & Wright, 2014).

All the language features were accessed from the URIEL knowledge base, queried with the lang2vec package (Littell et al., 2017). We correlated the pairwise distance metrics described above with the best-layer pairwise transfer scores obtained by $\text{InfoXLM}_{\text{large}}$, the best performing model for which encoding scores were available for all languages; significance testing accounted for false discovery rate given that we conducted multiple tests. **Supplementary Figure 12B** reports the results of the pairwise transfer. The correlational analyses relating the linguistic distance metrics with the transferability scores showed no meaningful pattern of association between language similarity and cross-lingual encoding transfer: indeed, transferability was not significantly associated with genetic ($r$ = 0.063, $p$ = n.s.), geographical ($r$ = 0.169, $p$ = n.s.), syntactic ($r$ = -0.202, $p$ = n.s.), phonological ($r$ = 0.083, $p$ = n.s.), or phonetic distance ($r$ = 0.275, $p$ = n.s.).

**Reliability of inter-language differences.** The absence of an association between encoding transferability and typological similarity could reflect a genuine attribute of the cross-lingual transfer effects we report: that linguistic form features do not strongly modulate the shared component in brain responses that is captured by the encoding models (consistent with the findings from Studies III and IV showing that the cross-lingual component is primarily semantic). However, another (not mutually exclusive) possibility is that variability in data quality across languages limits the ability to detect subtle typological effects.

To investigate this possibility, we examined the relationship between split-half reliability of the fMRI time series (i.e., the inter-participant correlation in each language; see **Figure 4A**) and cross-lingual transfer performance. We found that reliability was a significant predictor of a language's average transfer performance across all other languages ($r$ = 0.626, $p$ = 0.030; **Supplementary Figure 12C** shows this relationship at the level of pairwise transfer, where the average reliability of a language pair significantly predicted pairwise transfer: $r$ = 0.472, $p$ < 0.001). This result indicates that at least a substantial portion of the inter-language variability in transfer performance can be attributed to differences in data quality.

This observation also helps explain the zero and negative transfer values observed for certain language pairs in **Supplementary Figure 12B**. Of the 66 unique off-diagonal language pairs, 54 (82%) showed positive transfer. The 12 pairs with zero or negative transfer disproportionately involved the lowest-reliability languages: 6 of these pairs involved Marathi (the language with the lowest reliability, $r$ =

0.196), and 4 involved Dutch (the third-lowest, $r$ = 0.209). This pattern suggests that noise in the fMRI data is the most likely explanation for the low transfer scores observed in specific language pairs, and not absence of shared linguistic representations for specific language pairs.

**Within- versus across-language encoding.** Despite the prominent role of data quality in determining inter-language differences in encoding performance, we do find the expected pattern whereby within-language encoding outperforms across-language transfer. **Supplementary Figure 12D** compares, for each language, the within-language encoding performance (the diagonal of the pairwise transfer matrix in **Supplementary Figure 12B**) with the average across-language transfer (the off-diagonal mean). For 11 of the 12 languages, within-language performance exceeded across-language performance. The sole exception was Romanian (within-language $r$ = 0.060; across-language $r$ = 0.164). A permutation test confirmed that observing only 1 out of 12 reversals is significantly fewer than expected under random assignment of *within* versus *across* labels ($p$ = 0.002). This result confirms that the single-language transfer scores, while noisy, do carry some signal, and that the within-language encoding models capture some language-specific information beyond the shared cross-lingual component.

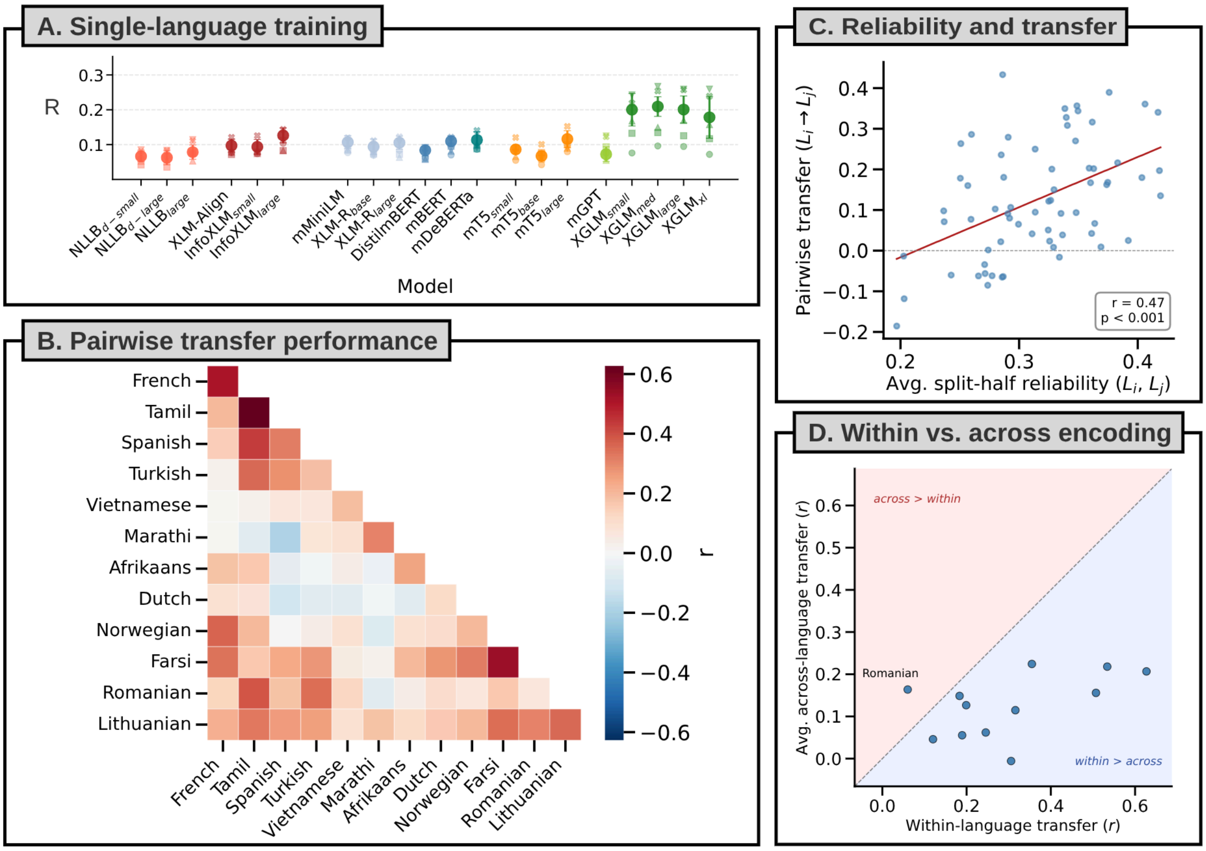


**Supplementary Figure 12:** *A.* Average transfer performance across models in the single-language training setup. Each dot indicates the best-layer performance of an individual model, averaged across languages. Error bars indicate the standard error across languages. Despite reduced training data, the relative ranking of model families mirrors the main results (cf. Figure 2A, bottom). *B.* Pairwise transferability heatmap for InfoXLMlarge. Values reflect the correlation between predicted and observed responses for each language pair, averaged across directions. None of the tested typological similarity metrics (syntactic, geographic, phonological, genetic, inventory, featural) significantly predicted transfer after correction for multiple comparisons. *C.* Relationship between pairwise data reliability and pairwise cross-lingual transfer for InfoXLM$_{large}$. Each dot represents a language pair (N = 66); the *x*-axis shows the average split-half reliability of the two languages, and the *y*-axis shows the pairwise transfer score. Reliability is significantly associated with transfer, suggesting that variability in data quality accounts for a substantial portion of inter-language differences in transfer performance. *D.* Within-language versus average across-language encoding performance for InfoXLM$_{large}$. Each dot represents one language. The dashed line indicates same within- and across-language performance; points below the line indicate the expected pattern where within-language encoding performance is higher than across-language performance. Romanian is the only language falling above the diagonal.

### S15. Language-level analyses

In the paper, we showed that the models' next-word prediction abilities accounted for inter-model differences in encoding performance (Results, "Next-word prediction ability explains inter-model differences in within-language encoding performance"). To assess whether those features additionally predicted differences in encoding performance across languages, we repeated those analyses following an identical procedure, but averaging across models instead of across languages. With this approach, for each language $l_i$, we obtained one single datapoint for encoding performance and perplexity (next-word prediction ability) by averaging those values across the models that supported $l_i$ (20 for French, Tamil, Spanish, Turkish, Vietnamese; 16 for Marathi, Afrikaans, Farsi, Romanian, Lithuanian; 15 for Dutch and Norwegian).

Differences in encoding performance across languages could not be reliably predicted by perplexity (WITHIN: $r = -0.0655$, $p$ = n.s.; ACROSS: $r = -0.1105$, $p$ = n.s.). We attribute this null result to the noise in the encoding scores obtained in each individual language, stemming from the different reliability of brain data for specific languages and the relatively low number of responses in each language (see also S14 and **Supplementary Figure 12C**).

### S16. Language resources

Between Study I and Study II, our experiments considered a total of 22 languages. In this section, we present an overview of the resources available for those languages, both in the context of NLP (focusing on the availability of textual data) and neuroscience research (focusing on the number of publications in language neuroscience). For the former, we considered as an index of resource availability the number of tokens in a given language, using the mC4 corpus as a reference (Xue et al., 2021). For the latter, we considered the estimates provided by Malik-Moraleda, Ayyash, et al. (2022). We report this information in **Supplementary Table 1**, which shows that our language sample is highly variable in terms of both NLP resources (with languages contributing from 2B up to 713B tokens to mC4) and coverage in neuroscience research, encompassing both understudied (<10 papers; 8 languages), somewhat studied (>10 but <100 papers; 7 languages), and well-studied languages (>100 papers; 6 languages).

Not surprisingly, we found that languages that are underrepresented in the language technologies tend to be understudied in language neuroscience as well (Spearman's $r = 0.50$, $p = 0.0201$), highlighting a systemic imbalance in the representation of languages across fields.

| Language | Study | Tokens (B) | fMRI studies | Language | Study | Tokens (B) | fMRI studies |
|---|---|---|---|---|---|---|---|
| English | N/A | 2733 | Well studied | Romanian | I | 52 | Understudied |
| French | I | 318 | Well studied | Lithuanian | I | 11 | Understudied |
| Tamil | I | 3 | Understudied | Hindi | II | 24 | Somewhat studied |
| Spanish | I | 433 | Well studied | Korean | II | 26 | Well studied |
| Turkish | I | 71 | Understudied | Arabic | II | 57 | Somewhat studied |
| Vietnamese | I | 116 | Understudied | Polish | II | 130 | Somewhat studied |
| Marathi | I | 14 | Understudied | Russian | II | 713 | Somewhat studied |
| Afrikaans | I | 2 | Understudied | Portuguese | II | 146 | Somewhat studied |
| Dutch | I | 73 | Well studied | Mandarin | II | 39 | Well studied |
| Norwegian | I | 27 | Somewhat studied | German | II | 347 | Well studied |
| Farsi | I | 52 | Understudied | Italian | II | 162 | Somewhat studied |

**Supplementary Table 1:** Summary of the resource available for the 21 languages considered across Study I and Study II. The column "Tokens (B)" indicates the number of tokens (billions) in the target language in mC4, a large-scale multilingual corpus. The column "fMRI studies" reports how much the language in question had been studied with fMRI experiments until 2022, as indicated by Malik-Moraleda, Ayyash, et al. (2022) (well studied: >100 papers per language; somewhat studied: >10 but <100 papers; understudied: <10 papers) We also report information for English for reference, even though it was not included in our sample.

### S17. Form- and meaning-ablation in English

In the main text (Study IV), we report the effects of INLP-based meaning and form ablation on cross-lingual transfer, training the encoding models using the *Tuckute2024* dataset and testing them on the data that we collected for Study II. Here, we report the results of the same procedure evaluated within English, using 5-fold cross-validation on the *Tuckute2024* dataset.

Within English, both meaning ablation and form ablation substantially reduced encoding performance relative to the intact condition. Intact embeddings achieved a mean cross-validated encoding performance of $r$ = 0.431 (*SE* = 0.033). Ablating meaning-related information from the embeddings reduced encoding performance to $r$ = 0.156 (*SE* = 0.091), conserving 36.2% of the intact performance. Similarly, ablating form-related information from the embeddings led to an encoding performance of $r$ = 0.161 (*SE* = 0.084), conserving 37.4% of the intact performance. Both reductions were statistically significant (meaning-ablated vs. intact: $z = -14.47$, $p < 0.0001$; form-ablated vs. intact: $z = -14.22$, $p < 0.0001$). Critically, the two ablation conditions did not differ from each other ($z = -0.25$, *n.s.*). This indicates that meaning and form features have a comparable contribution to within-language brain encoding in English (**Supplementary Figure 13**).

This pattern is in contrast with the cross-lingual transfer results reported in Study IV, where meaning ablation produced a much larger reduction in performance than form ablation. Thus, form-related features appear to be largely language-specific and do not transfer well across the typologically diverse languages in our sample.

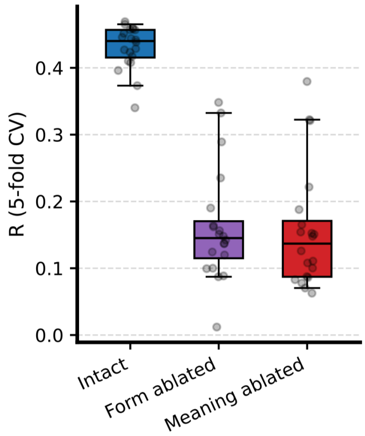


**Supplementary Figure 13.** Effect of INLP-based meaning and form ablation on within-English encoding performance (*Tuckute2024*, 5-fold cross-validation). Each dot represents one of the 20 MNNLMs. The boxplots show the distribution of mean cross-validated encoding scores across models for the three conditions: intact embeddings, form-ablated embeddings, and meaning-ablated embeddings. Both ablation conditions significantly reduce encoding performance relative to the intact condition (both $p < 0.0001$), but did not differ from each other ($p = 0.80$, *n.s.*).
